# Similar Fecal SCFA Patterns Despite Diverse Gut Microbiota in Polish and Japanese Children During the First Two Years of Life. The longitudinal, comparative, validated study

**DOI:** 10.1101/2025.10.23.684084

**Authors:** Igor Łoniewski, Mariusz Kaczmarczyk, Konrad Podsiadło, Danuta Cembrowska-Lech, Ulrike Löber, Robert Bücking, Karolina Skonieczna-Żydecka, Sofia K. Forslund, Beata Łoniewska

## Abstract

**Background:** Early-life gut microbiome assembly is shaped by environment, diet, and perinatal exposures. Whether population context shifts taxonomic trajectories and functional outputs similarly remains unclear.

**Methods:** We re-analyzed longitudinal infant cohorts from Poland (PL) and Japan (JP) across five time points (1 week, 1 month, 6 months, 1 year, 2 years). Amplicon sequence variants (ASVs) were processed uniformly. Cross-sectional alpha diversity (richness, evenness, Shannon, Simpson) was modeled with covariate adjustment (sex, antibiotics, diet). Beta diversity used Bray-Curtis distances (PCoA) and PERMANOVA (unadjusted/adjusted). Confounder-aware genus-level differential abundance identified features not reducible to covariates. Longitudinal genus trajectories and genus-SCFA (acetate, propionate, butyrate, isobutyrate) associations were tested using mixed-effects models. SCFAs were z-score–normalized to harmonize units across cohorts. Sensitivity analyses restricted PL to vaginally delivered infants. Finnish (FIN) and United States (US) cohorts were included for taxonomic validation.

**Results:** Evenness, Shannon, and Simpson were similar between PL and JP at most time points; richness was higher in PL at 1 week with trends at 6 months. Beta diversity showed a significant country effect at every time point except 1 month, robust to covariate adjustment and to restriction to 74 genera shared between PL and JP. Genus-level differential abundance yielded 33 features (26 enriched in PL, 7 in JP) without consistent cross-time recurrence. Longitudinally, most genus trajectories were cohort-specific; genus-SCFA associations were largely population-specific (few overlaps for butyrate/isobutyrate, none for acetate/propionate). Despite taxonomic and association differences, SCFA trajectories did not differ between cohorts after harmonization and adjustment. FIN/US validation supported the robustness of temporal taxonomic signals with few discordances.

**Conclusions:** Taxonomic profiles and genus-SCFA relationships diverge across populations, whereas core metabolic outputs (fecal SCFAs) follow a conserved, resilient trajectory in early life. This “divergent taxa, convergent function” pattern suggests early-life interventions should prioritize functional maturation over targeting specific taxa. Broader multi-omic studies integrating growth, immune, and neurodevelopmental outcomes are warranted.

## Introduction

Microbial colonization of the human gut begins soon after birth and undergoes dynamic transformations throughout early childhood, particularly during the first three years of life (Yatsunenko et al., 2012; Stewart et al., 2018; Roswall et al., 2021; Wernroth et al., 2022; Tsuruoka et al., 2024; Fahur Bottino et al., 2025). The infant intestinal microbiota is typically dominated by facultative anaerobic bacteria, such as *Enterobacteriaceae*, and exhibits low total number bacterial cells and diversity (Barker-Tejeda et al., 2024, Xiao and Zhao, 2023). Within a few days of life, the intestinal environment becomes more anaerobic, facilitating the proliferation of *Bifidobacterium*, which subsequently dominates the gut microbiota during the initial months of life due to its specialized metabolism of human milk oligosaccharides (Pucci et al., 2025; Samarra et al., 2025, Xiao and Zhao, 2023). The infant gut microbiota diversifies progressively during the first months of life, influenced by factors such as environmental exposures, contact with different microbes, and immune system maturation. The introduction of solid foods at approximately six months accelerates this diversification, leading to an adult-like microbiome dominated by *Firmicutes* and *Bacteroidetes* (Rodríguez et al., 2015; Stewart et al., 2018).

Research to date has mainly focused on assessing the impact of various environmental factors, such as mode of delivery, feeding, exposure to antibiotics, and even prenatal exposures to trace elements on the development of the microbiome (Bäckhed et al., 2015; Bokulich et al., 2016; Shao et al., 2019; Xiong et al., 2025). Geographical and cultural factors also markedly influence microbial colonization patterns, resulting in distinct microbiome trajectories across populations. The results of cross-sectional studies comparing microbiota in children from different geographical areas showed significant differences (Kuang et al., 2016; Ho et al., 2018; Fahur Bottino et al., 2025). The development of the early gut microbiota has immediate and long-term consequences for the health of the host and may be linked to later obesity (Dogra et al., 2015), risk of asthma (Arrieta et al., 2015; Stokholm et al., 2018), diabetes type 1 (Kostic et al., 2015; Stewart et al., 2018; Vatanen et al., 2018) and state of nutrition (Subramanian et al., 2014). In particular, the composition and function of early gut microbiota significantly impact immune system maturation, potentially affecting susceptibility to chronic inflammatory and metabolic diseases later in life (Hickman et al., 2024; Bui et al., 2025). Thus, understanding the intricacies of early-life microbiome establishment, influenced by maternal microbiota, feeding methods, environmental exposures, and geography, remains crucial for developing targeted interventions aimed at optimizing infant health trajectories.

Among the mechanisms responsible for the influence of the gut microbiota on host health, short-chain fatty acids (SCFAs) play a fundamental role (Louis et al., 2014; Koh et al., 2016; Morrison and Preston, 2016a; Ríos-Covián et al., 2016). SCFAs production depends primarily on the amount of fiber consumption and the composition and metabolic capacity of the microbiome (Dalile et al., 2019). *Clostridium*, *Eubacterium* and *Fusobacterium* (Bourriaud et al., 2005) are the major genera responsible for butyrate production, whilst the most intensive synthesis is performed by *Clostridium leptum*, *Roseburia* spp., *Fecalibacterium prausnitzii*, and *Coprococcus* spp (Guilloteau et al., 2010). Propionate production, conversely, is typically associated with the metabolic activities of Bacteroidetes and *Propionibacterium* (Salonen et al., 2014). Furthermore, SCFAs such as acetate can undergo conversion to propionate and butyrate through cross-feeding interactions involving various Firmicutes and Bacteroidetes representatives (Mahowald et al., 2009). These processes however are of individual nature and reflect the abundance of each major SCFAs producers (Czajkowska and Szponar, 2018). As elegantly summarized by others (Hu et al., 2018; Dalile et al., 2019; Deleu et al., 2021) the pivotal roles played by SCFAs in host physiology extend across multiple systems, including the regulation of carbohydrate metabolism (Portincasa et al., 2022), control of cholesterol synthesis and fatty acid metabolism (den Besten et al., 2013), regulation of energy intake (Chambers et al., 2015) and control of the homeostasis within the digestive system (van der Hee and Wells, 2021). SCFAs also possess anti-inflammatory and anti-cancer effects (Feitelson et al., 2023), regulate the nervous system (Lan et al., 2024) and have impact on immunomodulation (Ratajczak et al., 2019). Importantly, recent studies emphasize the immunomodulatory capacity of SCFAs, particularly in early life, where they shape adaptive immune responses and confer protective effects against inflammatory and allergic diseases (Bui et al., 2025; Samarra et al., 2025). SCFAs have been also shown to influence bacterial gene expression, which can affect microbial virulence and community dynamics (Lawhon et al., 2002; Nakanishi et al., 2009). These findings highlight the significance of SCFAs as critical intermediaries linking gut microbiome composition with host health outcomes, indicating the need for a deeper understanding of their production dynamics and biological functions.

Our research group has previously conducted several studies exploring the dynamics of gut microbiota development in cohort of healthy children during the first two years of life. We demonstrated that the gut microbiota composition, small intestinal paracellular permeability, and immune system-related markers undergo dynamic changes influenced by multiple perinatal factors (Kaczmarczyk et al., 2021). Additionally, we observed that fecal short-chain fatty acid (SCFA) concentrations fluctuate significantly during the first year of life before stabilizing. Furthermore, various maternal and fetal factors such as antibiotic use during pregnancy, delivery method, feeding practices, maternal body weight (both pre-pregnancy and gestational changes), as well as the infant’s sex and birth weight, have diverse impacts on SCFAs excretion in the stool (Łoniewska et al., 2023). In a comprehensive analysis including 41 studies (*n* = 2,457 SCFAs analyses), we identified feeding mode as a crucial determinant of early-life SCFA concentrations, showing increased fecal SCFA levels in artificially fed infants, whereas breastfed infants exhibited more dynamic SCFA fluctuations at different developmental stages (Łoniewski et al., 2022). However, despite significant evidence indicating geographical differences in microbiota composition, there is currently a notable gap in comparative longitudinal studies linking microbiota composition and metabolic function, specifically SCFA production, across distinct populations. Therefore, we conducted a comparative re-analysis aiming to elucidate differences in gut microbiota composition and associated fecal SCFA profiles between Polish infants and infants from other geographic regions during the first two years of life. Using publicly available datasets, we also tried to identify potential microbiota-SCFA associations that may vary geographically, thus contributing novel insights into the interplay between microbial ecology and metabolic function in early-life development.

## Methodology

### Studied cohorts

#### Polish cohort

The Polish cohort (PL) consisted of newborns born at the Department of Obstetrics, Gynecology and Neonatology at the Pomeranian Medical University/Independent Public Clinical Hospital No. 2 in Szczecin and was sampled between March 2015 and April 2016. This cohort has been previously described in detail in Łoniewska et al. (2019), Łoniewska et al. (2020), and Kaczmarczyk et al. (2021), referred to as PMU cohort in the latter. A total of 107 stool samples were collected at predetermined intervals: meconium (birth, N = 10), 7 days (1 week, N = 23), 1 month (N = 20), 6 months (N = 20), 1 year (N = 21), and 2 years (N = 13). Stool collection and DNA extraction were previously described in Kaczmarczyk et al. (2021). Briefly, stool samples were collected from diapers using a biological material collection kit (Stool Sample Application System (SAS); Immundiagnostik, Bensheim, Germany). The stool samples were then frozen at − 20 °C until metagenomic analyses were conducted. Microbiome DNA extraction was performed using the Genomic Mini AX Bacteria + Spin and Genomic Mini AX Soil Spin kits (A&A Biotechnology, Gdynia, Poland) following the manufacturer’s protocol. Metagenomic libraries of the V3-V4 region of the 16S rRNA gene were constructed and sequenced on the MiSeq platform (paired-end 2 × 250 bp) using V2 chemistry from Illumina (Illumina, San Diego, CA, USA).

#### Data comparing the microbial composition of infant gut microbiota across diverse countries - Database search

Two public databases, the Sequence Read Archive (SRA) and Microbiome DB (https://microbiomedb.org/mbio/app), were searched for relevant datasets of clinically healthy infants using the following criteria: infant gut microbiota with 16S rRNA gene sequencing for microbial community analysis; only publicly available datasets with a minimum total sample size of 50, age range of infants within the 0 - 24 months and at least four time points to allow comparisons with the PL cohort. As of 2021, three cohorts that met these criteria were included: the Finnish cohort (FIN), the Japanese cohort (JP), and the ECAM United States cohort (US). Detailed characteristics of the newborn cohort included in the study and obtained from the public resources are provided in Supplementary Table 1 and 2. An initial screening of metadata revealed the presence of SCFAs data only in the JP. Given the role of SCFAs in host metabolic and immunological regulation, we focused on the SCFAs levels and their possible association with gut microbiota. The remaining cohorts (FIN and US) were used for validating methods used for the analysis, as well as microbial compositions and longitudinal dynamics, observed in the JP and PL cohorts.

#### Sample selection algorithm and time points harmonization across cohorts

To harmonize cohort data by time point, one sample per individual was selected from the FIN, US and JP datasets to match each time point defined in the PL cohort. In cases where no exact-match sample was available for a given timepoint, a ±10% age tolerance was applied (e.g. for the 1 month timepoint, a sample was considered eligible if it fell within the timeframe of 27 to 33 days). If multiple samples met the specified criteria for an individual, a random sample was chosen. The harmonized timepoints, and the number of infants per time point for each cohort are provided in Supplementary Table 2. Age bin ranges for JP samples (in days) were: 7–7.5 (1st week), 29 (1 month), 157–183 (6 months), 356–364 (1st year), and 690–735 (2nd year). In the PL cohort, bins included the first stool and the 1st week (7 days, collected at the hospital), 30 and 180 days ±3 days (1st and 6th month), and 365 and 730 days ±7 days (1st and 2nd year). To verify that subsampling of the JP cohort did not affect SCFA dynamics, SCFA levels from the full cohort and the subset of selected individuals were visualized and compared (Supplementary Figure 1). Full metadata for the JP cohort and the subsample used in this study are available in Supplementary Table 3.

#### Raw sequences processing, taxonomic assignment and data processing

An initial quality check was performed using FastQC (v0.12.1) and summarized with MultiQC (v1.23). After this step, the sequencing data was processed through the LotuS2 pipeline (Özkurt et al., 2022). Quality filtering and dereplication were carried out using the sdm module tailored for the MiSeq or HiSeq Illumina platform, with a slightly reduced minimum sequence length. Within LotuS2, sequence clustering into ASVs (Amplicon Sequence Variants) was achieved using the DADA2 algorithm (Callahan et al., 2016). To remove chimeric sequences, both reference-based and *de novo* detection methods were applied using the RDP reference database (rdp_gold.fa) with UCHIME3 (Edgar, 2016). By default, ASVs were aligned against the phiX genome using Minimap2 (Li, 2018), and any sequences showing significant similarity were excluded. Reads that matched the human reference genome (Homo_sapiens.GRCh38.dna.toplevel.fa; release 2024-02-13) were also filtered out to eliminate host contamination.

The dataset was further refined using the LULU algorithm (Frøslev et al., 2017), which curated sequence clusters. The remaining ASVs were aligned using Lambda v3.0.0 (Hauswedell et al., 2014) against the SILVA 138.1 SSU database (Yilmaz et al., 2014) to assign taxonomy through LotuS2’s default Least Common Ancestor (LCA) method. Sequences identified as Eukaryota, Archaea, mitochondria, or chloroplasts were retained in the dataset for alpha-diversity analysis.

Four alpha-diversity metrics (richness, evenness, Simpson index, Shannon index) were determined for each project using a common rarefaction depth of 5,312 and the *get.mean.diversity* function from the *rtk* (version 0.2.6.1) R library. Additionally, alpha-diversity metrics were re-calculated for ASVs mapped to shared genera (N = 74) between the PL and JP cohorts and a rarefaction depth of 5,062. For taxonomic-based analyses, sequences identified as Eukaryota, Archaea, mitochondria or chloroplasts were removed from each dataset after rarefaction. ASVs with incomplete taxonomic classification were retained and labeled according to their last confidently assigned taxonomic level. This filtering resulted in the following number of genera per cohort: 491 (PL), 98 (JP), 335 (US), 161 (FIN, HiSeq), and 247 (FIN, MiSeq). The resulting genus-level taxonomic tables from each dataset were combined yielding a total of 661 unique genera across all cohorts. A 10% prevalence filtering (i.e., presence in at least 10% of samples with one or more assigned reads) was applied resulting in 89 genera. For analyses limited to the PL and JP cohorts, the respective genus-level tables were merged separately, resulting in a total of 501 unique genera. After applying the 10% prevalence filter, this was reduced to 97 genera overall and 95 shared between both cohorts. Compositional differences were assessed using the Bray–Curtis (BC) distance at the genus level. For visualization, Principal Coordinates Analysis (PCoA) was used to display sample relationships based on compositional dissimilarity. To test for differences in community structure between groups, Permutational Analysis of Variance (PERMANOVA) using the adonis2 function in the vegan R package (version 2.6-10). The by = “margin” was specified, which reports marginal effects for each term (adjusting for the others). BC distances were calculated using the full set of samples, while both PCoA plots and PERMANOVA tests (9,999 permutations) were performed on time point - specific subsets of the distance matrix.

Cross-sectional genus-level differential abundance analysis was performed using metadeconfoundR package (version 1.0.2) in which univariate associations between taxa and cohort (PL vs JP) are checked for confounding effects and one of the four statuses are returned accordingly. The statuses are the following: OK_nc (no covariates - cohort is the only significantly associated covariate), C (confounded), AD (ambiguously confounded - another covariate is associated with a taxon, but neither association is significantly stronger than the other), OK_d (doubtful - another covariate is associated with a taxon, but the signal can be reduced to this covariate, confidence interval includes 0), and OK_sd (another covariate is associated with a taxon, but the signal can be reduced to this covariate). To identify taxa that changed significantly over time while accounting for covariates we used LongDat R package (version 1.1.3, Chen et al., 2023), treating time as a discrete variable. The analysis was performed separately for PL and JP cohorts, for both consecutive and non-consecutive time points. Consecutive comparisons included the following intervals: 1 week vs. 1month, 1 month vs. 6 months, 6 months vs. 1 year, 1 year vs. 2 years. For non-consecutive, we used 1 week, 1 month, and 6 months as static reference, and tested against all later time points not already included in the consecutive analyses. OK_nc (no covariates - cohort is the only significantly associated covariate), OK_d (doubtful - there is an effect of time and there is no potential covariate, however the confidence interval of the time estimate in the model test includes zero), OK_nrc (there are potential covariates, but the effect of time is independent of covariates), EC (entangled with covariates - there are potential covariates, and the effect can result from time or covariates), RC (there is an effect of time, but it can be reduced to the covariate). Alpha-diversity, PERMANOVA, cross-sectional differential abundance analysis (DAA) and longitudinal analyses were adjusted for sex, diet, and use of antibiotics (ATB). As the Japanese cohort consisted exclusively of vaginally-delivered newborns, a sensitivity analysis was performed by repeating the analyses in a subset of the PL cohort limited to vaginally delivered newborns.

#### SCFAs Profile Analysis and Cross-Cohort Comparison

SCFAs analysis was conducted with methods described previously in Łoniewska et al. (2023) and Tsukuda et al. (2021). Cross-cohort comparison of SCFAs levels, given their expression on distinct scales (mM and nM/mg in Japanese and Polish, respectively), was addressed through z-score normalization. Trends in normalized SCFAs levels between the Polish and Japanese cohorts were then analyzed using mixed-effect models (lme4 package, version 1.1-35.3) adjusted also for sex, diet, and ATB, and also in a subset of vaginally born Polish newborns.

## Results

### Basic characteristics of the groups

Samples from newborns were collected at multiple time points, and metadata including delivery type, diet type, antibiotic usage, gender, and birth weight were recorded and tabulated (**Table 1**). The Polish cohort also included data on birth weight.

**Table 1.** Clinical characteristics of newborns by Polish and Japanese cohort.

| Timepoint | Project | Delivery:<br>Vaginal,<br>N (%) | Diet type:<br>Breastfeeding,<br>N (%) | Antibiotics<br>treatment<br>history, N (%) | Sex:<br>Female,<br>N (%) | Birth weight<br>(kg), M (SD) |
| --- | --- | --- | --- | --- | --- | --- |
| Birth | PL | 3 (33%) | 9 (100%) | 0 (0%) | 4 (44%) | 3.5 (0.6) |
|  | JP | - | - | - | - | - |
| 1st week | PL | 12 (52%) | 21 (91%) | 1 (4%) | 10 (43%) | 3.3 (0.5) |
|  | JP | 12 (100%) | 7 (58%) | 1 (8%) | 5 (42%) | - |
| 1 month | PL | 9 (45%) | 15 (75%) | 1 (5%) | 11 (55%) | 3.3 (0.5) |
|  | JP | 12 (100%) | 10 (83%) | 0 (0%) | 5 (42%) | - |
| 6 months | PL | 8 (40%) | 10 (50%) | 7 (35%) | 9 (45%) | 3.3 (0.5) |
|  | JP | 12 (100%) | 2 (17%) | 0 (%) | 5 (42%) | - |
| 1 year | PL | 9 (43%) | 4 (19%) | 12 (57%) | 10 (48%) | 3.3 (0.5) |
|  | JP | 12 (100%) | 0 (0%) | 1 (8%) | 5 (42%) | - |
| 2 years | PL | 7 (54%) | 1 (8%) | 10 (77%) | 8 61%) | 3.3 (0.5) |
|  | JP | 12 (100%) | 0(0%) | 1 (8%) | 5 (42%) | - |

PL- Polish cohort, JP- Japanese cohort, M - mean, SD - standard deviation

### Cross-sectional microbiome diversity and abundance analyses in Japanese vs. Polish newborns

Cross-sectional comparisons between the PL and JP cohorts were conducted using linear models that included antibiotic exposure (ATB), sex, and diet as covariates, except at the 2-year timepoint, where none of the newborns were breastfed. Four alpha diversity indices-richness, evenness, Shannon, and Simpson - were obtained from ASVs (after removing samples with a total count ≤ 5,000 and rarefying to a common depth of 5,312, and were assessed at each time point. Detailed statistics on ASVs per sample and per feature are presented in Supplementary Table 4. No significant differences were observed between the PL and JP cohorts in terms of evenness, Shannon, and Simpson indices (Figure 1A-C), although a trend for a higher diversity in the PL cohort was noted at 6 months (evenness: FDR-adjusted p-value [Q] = 0.059; Shannon: Q = 0.054). For richness, a similar trend was observed at 6 months (Q = 0.059), and a significant difference was detected at 1 week (Q = 0.002; Figure 1E, left panel). Full results of this cross-sectional analysis of the alpha diversity metrics are provided in the Supplementary Table 5 and Supplementary Figure 2. Although only richness at 1 week differed significantly between the PL and JP cohorts (Figure 1E), we applied a resampling approach with fixed and equal sample sizes - ranging from 25 to 50 - and with 100, 300, and 500 iterations for both cohorts (PL: N = 105; JP: N = 59), in order to assess whether the sample size imbalance could affect the total number of ASVs per cohort and thereby influence alpha diversity metrics. The differences between the Poland and Japan cohorts in terms of the total ASV number, ASVs mapped to taxonomy at the genus level, and proportions of assigned/unassigned taxa to ASVs at the genus level were relatively consistent across sample sizes, thereby implying that cohort-level differences are not driven by sampling depth or sample size (Supplementary Figure 3).

**Figure 1.**
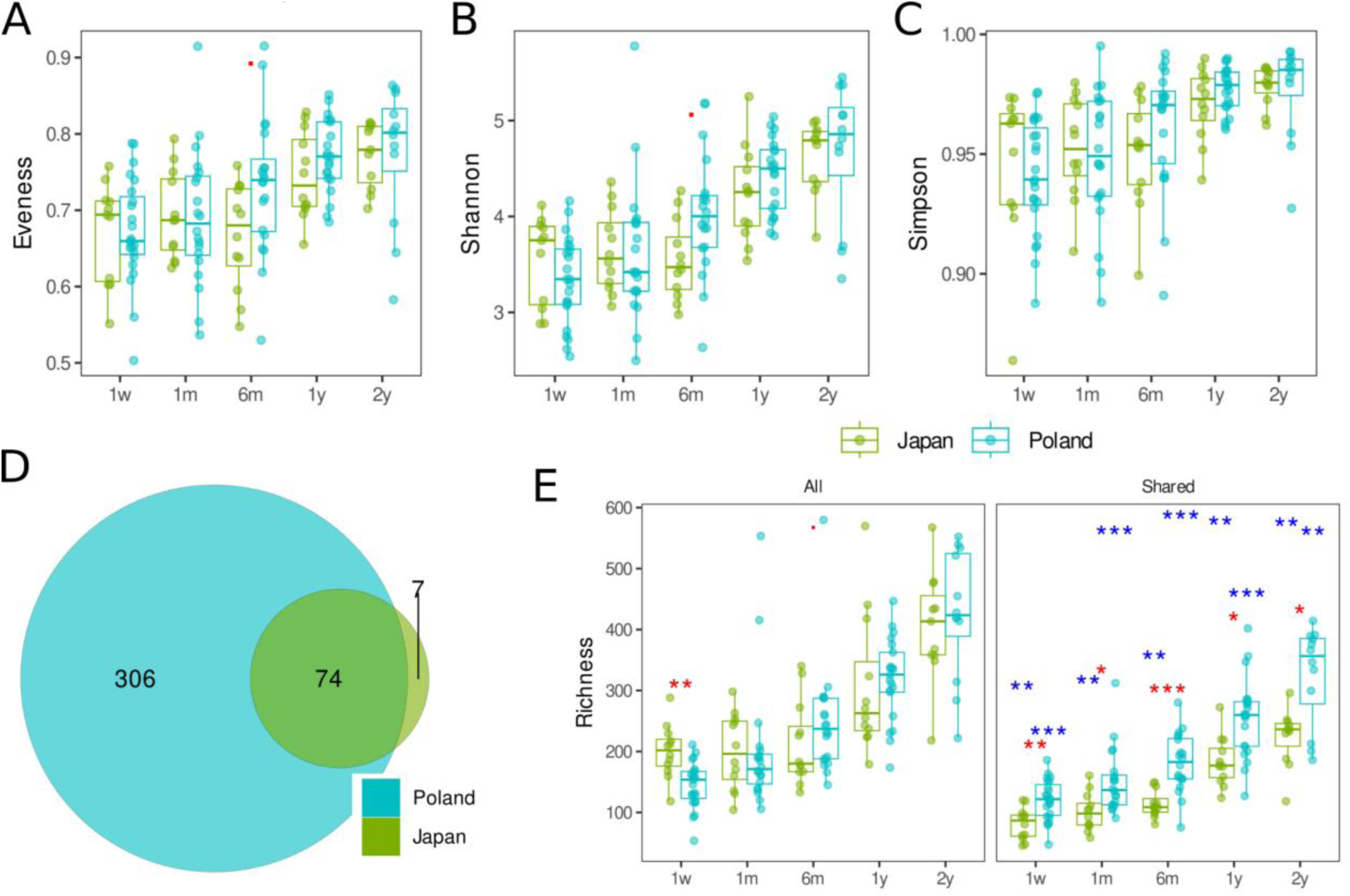
**Alpha diversity comparisons between Poland and Japan cohorts – all ASVs and ASVs from shared genera.** A-C: Alpha diversity metrics calculated using all ASVs: (A) Evenness, (B) Shannon index, and (C) Simpson index, across five time points for PL and JP cohorts, D: Venn diagram illustrating the number of genera unique to each cohort and those shared (N = 74) between PL and JP, E: Comparison of richness estimates across cohorts and analysis scopes. Left panel: richness calculated using all ASVs (non-restricted). Right panel: richness based on ASVs mapped to shared genera (restricted analysis). Red asterisks denote statistically significant differences between PL and JP cohorts at individual time points (Q < 0.05). Blue asterisks indicate significant differences between non-restricted and restricted diversity metrics at the same time point (paired Wilcoxon test, Q < 0.05). 0 - 0.001 ’***’, 0.001 - 0.01 ’**’, 0.01 - 0.05 ’*’, 0.05 - 0.1 ’.’, 0.1 - 1.0 (No symbol) 1w - 1 week of life, 1m - 1 month, 6m - 6 months, 1y - 1 year, 2y - 2 years.

To assess whether observed diversity patterns were driven by the presence of cohort- specific taxa, we recalculated alpha diversity metrics using only the subset of ASVs mapped to shared genera (N = 74, Figure 1D) between the PL and JP cohorts. By mapping to common genera we intended to reduce 16S region-dependent artifacts (ASVs defined differently but referring to similar organisms) thereby measuring how diversely the same genera are populated in PL vs. JP. After we restricted the analysis to ASVs mapped to a set of 74 genera shared between cohorts, alpha diversity metrics became overall significantly lower than non- restricted metrics (paired Wilcoxon test, blue asterisks in Figure 1 E, right panel), indicating the influence of unassigned and cohort-specific ASVs on diversity inflation. Importantly, the restricted analysis revealed significant differences between PL and JP samples (red asterisks, 1 week: Q = 0.002, 1 month: Q = 0.025, 6 months: Q = 0.00009, 1 year: Q = 0.033, 2 years: Q = 0.017, Figure 1 E, right panel). Full results of this restricted cross-sectional analysis of the alpha diversity metrics are provided in the Supplementary Table 6 and Supplementary Figure 4, while paired non-restricted and restricted in Supplementary Table 7.

Beta-diversity was based on Bray-Curtis (BC) distances that was calculated on the full set of all samples (all timepoints) and then used to ordinate samples in a two-dimensional space using Principal Coordinate Analysis (PCoA) separately for each timepoint (1w, 1m, 6m, 1y, and 2y, Figure 2A-B). The time point specific subset of the BC distance matrix obtained from all samples were subjected to PERMANOVA for association analysis including country (non-adjusted analysis) and covariate-adjusted analysis including Country, sex, antibiotic use, and type of diet (except for the 2 years time point where all newborns were on the same diet). Considering both shared and unique genera we observed a significant (Q < 0.1) influence (also after adjusting for covariates) of country on the microbial composition in all time points (Figure 2A).

**Figure 2.**
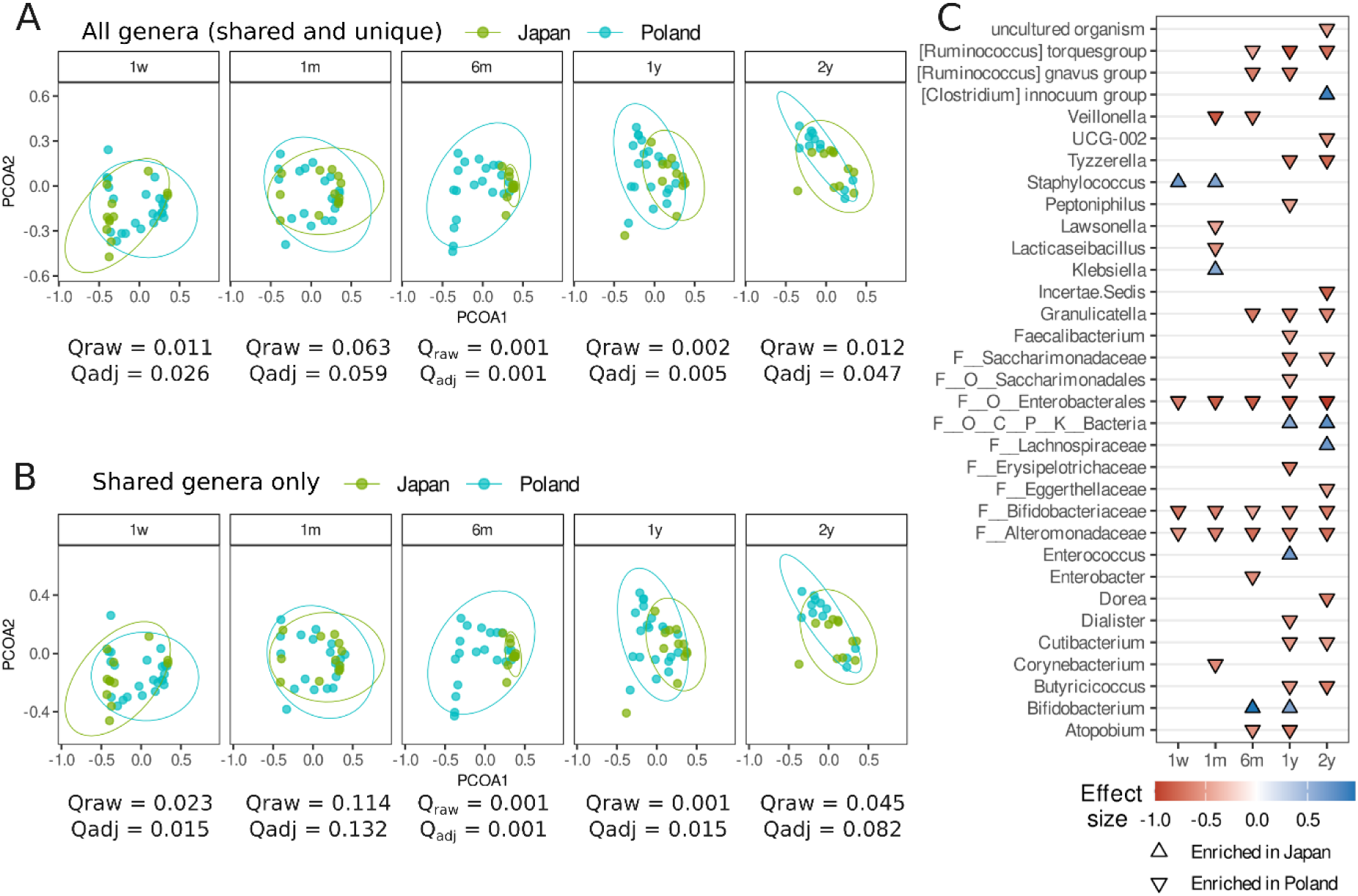
**Genus-level beta-diversity by country and differential abundance analysis adjusted for confounders** A, B - PCOA - upper panel - based on Bray-Curtis distances for all genera (10% prevalence filter), lower panel - based on Bray-Curtis distances for 73 shared genera (10% prevalence filter), 1w - 1 week, 1m - 1 month, 6m - 6 months, 1y - 1 year, 2y - 2 years, 73 shared genera, Praw - PERMANOVA P value for comparison PL and JP, Padj - PERMANOVA P value for comparison PL and JP adjusted for ATB and delivery; C - deconfounding analysis - separately for each time point, accounting for sex, diet, ATB - only deconfounded features are shown, when a prefix is present (e.g. F) it indicates carry-over from the previous known taxa, positive association - enriched in JP cohort, negative association - enriched in PL cohort, for all genera, Effect size - Cliff’s Delta, only features with OK_nc and OK_sd statuses are reported: OK_nc (No other covariates): Country is the only covariate associated with the feature), OK_sd (Strictly deconfounded): another covariate is associated with the feature, but the signal can not be reduced to this covariate)

To assess whether the observed community-level differences were driven primarily by the overall genus composition or by genera shared between countries, we repeated the BC-based PCoA and PERMANOVA analyses on a subset of the data restricted to 74 (Figure 2B) genera shared between the PL and JP cohorts. The beta-diversity distance matrix was recalculated using only these shared genera, and all subsequent ordinations and PERMANOVA models (including unadjusted and covariate-adjusted comparisons) were re-run. The results were highly consistent with those from the full-genus dataset (Figure 2A). Specifically, the time-point specific patterns and significance of group-level differences (Q < 0.1) (including the lack of significance at 1 month) remained unchanged. This suggests that the core microbial structure shared between populations, rather than cohort-specific genera, largely explains the beta-diversity patterns and the differences between cohorts. The complete PERMANOVA results across all scenarios (full set, shared genera, unadjusted, and covariate-adjusted) are provided in Supplementary Table 8.

To complement the beta-diversity analyses and identify specific microbial features potentially driving the observed community-level differences, we next carried out cross-sectional (within–time-point) comparisons at the genus level using a confounder-aware approach - that is, by detecting differentially abundant genera between the two cohorts where the association signal cannot be attributed to the covariate. A cuneiform plot depicting features that were strictly deconfounded, i.e. their association signal could not be reduced to the included covariates (sex, antibiotic use, type of diet), or for which Country was the only significant associated covariate, is presented in Figure 2C. We did not observe genera consistently differing between cohorts across multiple time points (except for features not classified at the genus level and with a taxonomy carried-over from higher ranks, e.g. F_Bifidibacteriaceae that were consistently enriched in the PL cohort over all time points). Among 33 differentially abundant genera (Q < 0.1), the majority (N = 26, 78.8%) was enriched in the PL cohort, and 7 genera (21.2%) were more abundant in the JP cohort (classified to a genus level: *Staphylococcus* - 1 week, Cliff’s delta (D) = 0.72, Q = 0.016, 1 month, D = 0.62, Q = 0.040; *Klebsiella* - 1 month, D = 0.59, Q = 0.040, *Bifidobacterium* - 6 months, D = 0.97, Q = 0.0006, 1 year, D = 0.62, Q = 0.018, *Enterococcus* - 1 year, D = 0.69, Q = 0.011, *[Clostridium] innocuum* group - 2 years, D = 0.89, Q = 0.009). Complete list of identified differentially abundant features is provided in Supplementary Table 9.

As all Japanese infants were delivered vaginally, we did not consider this variable an independent factor in between-cohort comparisons. Therefore, sensitivity analyses were performed by restricting the Polish cohort to vaginally delivered infants. In the non-restricted ASV analysis, results remained consistent between the full cohort and the subset of vaginally born infants (Supplementary Figure 2, Supplementary Table 5 vs. Supplementary Figure 5, Supplementary Table 10). Similarly, in the restricted ASV analysis (mapped to shared genera), the findings were consistent between the full cohort and the vaginally born subset (Supplementary Figure 4, Supplementary Table 6 vs. Supplementary Figure 6, Supplementary Table 11). However, as in the full cohort, the paired restricted and non-restricted alpha-diversity analyses (stratified by time point and cohort) showed significant differences between the two approaches (Supplementary Table 12). PERMANOVA conducted within the subset of vaginally born newborns yielded results largely consistent with those from the full cohorts (Supplementary Table 13). In the differential abundance analysis (DAA), 26 differentially abundant genera were identified, compared to 33 in the full cohorts (Figure 2C). Of these, 25 genera overlapped between the two analyses, while the genus group *Burkholderia*– *Caballeronia*–*Paraburkholderia* emerged uniquely as differentially abundant in the vaginally born subset. Notably, some genera that were differentially abundant in the full cohort - *Cutibacterium*, *Dorea*, *Faecalibacterium*, *Staphylococcus*, and *UCG-002* - were no longer significant when the analysis was restricted to vaginally born infants (Supplementary Table 14).

### Longitudinal SCFA dynamics in Japanese and Polish cohorts: involvement of microbial features

To explore gut microbial dynamics over time, we applied a covariate-aware genus-level analysis, adjusting for sex, antibiotic use, and diet. Analyses were focused on genera shared across both populations, and filtered at a prevalence of ≥10%. Sensitivity analyses were also conducted by restricting the Polish cohort to vaginally delivered infants. Results for consecutive time intervals - 1 week to 1 month, 1 month to 6 months, 6 months to 1 year, and 1 to 2 years are shown in Figure 3 (Panels A and B: full cohorts; Panels C and D: vaginally born subset). Results for additional intervals, including static time point comparisons starting at 1 week, 1 month, and 6 months, are presented in Supplementary Figures 7-9 (full cohorts) and Supplementary Figures 10-12 (vaginally born subset). In the full cohorts, the Polish group exhibited 35 significant changes in genus abundance, of which 3 were either entangled with or reducible to covariates. The Japanese group showed 21 significant changes, all independent of covariates (Figure 3A). However, only four genera were shared between cohorts: *Staphylococcus* (1 week - 1 month interval), *Romboutsia*, *Intestinibacter* (reducible to covariates in PL), and *Haemophilus* (6 months - 1 year interval, Figure 3B, Supplementary Table 15). A similar pattern of cohort-specific dynamics was observed when the PL was limited to the vaginally delivered infants. The PL cohort showed 17 significant changes (3 of which were entangled with or reducible to covariates, Figure 3C). The same four genera were again shared between cohorts: *Staphylococcus* (1 week - 1 month and 1 month - 6 months intervals), *Romboutsia* (entangled with covariates in PL), *Intestinibacter*, and *Haemophilus* (6 months - 1 year, Figure 3D, Supplementary Table 15).

**Figure 3.**
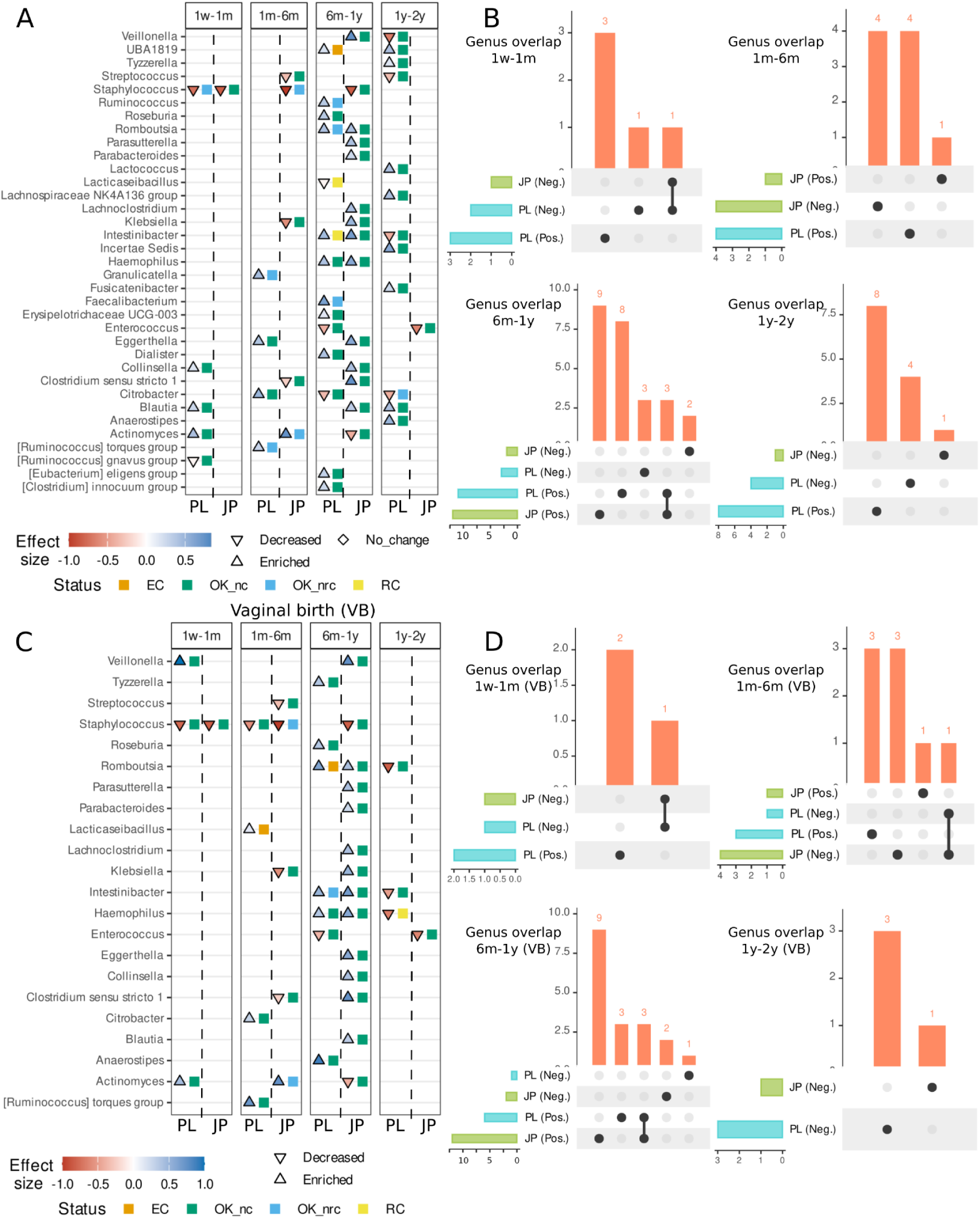
**Country-specific significant changes in genus-level abundance between adjacent (consecutive) time points and overlap of genus-level signals in full and vaginally-born cohorts** A, C - cuneiform plots displaying features whose signals were not classified as “NS” (non-significant) in the full cohorts (A) and in vaginally delivered newborns (VB) (C) - the direction of effects across the respective time intervals are shown. A triangle pointing downward indicates a decrease in abundance between time points, while Enriched denotes an increase. Effect sizes are represented by Cliff’s delta. Status labels definitions: EC (Entangled with covariate) - there are potential covariates, and it is not possible to conclude whether the effect is resulted from time or covariates; RC (Reducible to covariate) - there is an effect of time, but it can be reduced to the covariate effects; OK_nc (OK, no covariate) - there is an effect of time and there is no potential covariate; and OK_nrc (OK, not reducible to covariate) - there are potential covariates, but there is an effect of time and it is independent of those of covariates. B, D - UpSet plots illustrating the overlap of significant genera between countries. The intersection size bar plot (vertical) indicates the number of genera shared across different country - effect combinations, while the set size (horizontal) bars represent the total number of genera per group: PL (Neg.) - Polish cohort, decrease in abundance between time points, PL (Pos.) - Polish cohort, increase in abundance between time points, JP (Neg.) - Japanese cohort, decrease in abundance between time points, JP (Pos.) - Japanese cohort, increase in abundance between time points; Only individuals with complete samples at the compared time points were included.

Additional genera were identified as shared between countries when other time intervals were considered. For example, *Lachnoclostridium* and *Clostridium sensu stricto 1* were shared in the 1 week - 1 year interval in both the full and vaginally delivered cohorts (Supplementary Figures 7 and 10, Supplementary Table 15). *Lactococcus* was shared in the 6 months - 2 years interval in the full cohorts (Supplementary Figure 9, Supplementary Table 15), while *Flavonifractor* was shared in the same interval in the vaginally delivered cohort (Supplementary Figure 12, Supplementary Table 15). Overall, genera with a consistent temporal trend across cohorts accounted for only a minority of the significantly altered genera across the evaluated time points in full cohorts (from 7.7% to 20.0%, median 9.1%) and in the vaginally delivered (from 6.3% to 33.3%, median 11.5%) (Supplementary Table 15), supporting a cohort-specific pattern of time-dependent genus-level trajectories during the first two years of life.

We next investigated potential links between microbial dynamics and SCFA metabolism through a longitudinal taxonomic analysis of the JP and PL cohorts. Linear mixed-effects models between microbial taxa and SCFAs were applied, adjusting for covariates including sex, antibiotic use, and diet. Figure 4A presents genera significantly associated with four SCFAs (likelihood ratio test [LRT] P < 0.05) in the full Polish and Japanese cohorts, including both vaginally (VB) and cesarean-delivered (CS) participants. In total, 23 significant (LRT P < 0.05) genus SCFA associations were identified in the Polish cohort: acetate (N = 5), propionate (N = 8), butyrate (N = 7), and isobutyrate (N = 3), and 32 associations in the Japanese cohort: acetate (N = 3), propionate (N = 8), butyrate (N = 15), and isobutyrate (N = 6). However, these genus - SCFA associations differed substantially between the two cohorts. The UpSet plot (Figure 4B) shows that only two genera are associated with butyrate in the PL and JP cohort: *[Clostridium] innocuum* group (associated in opposite directions) and *Clostridium sensu stricto 1* (associated in the same direction). For isobutyrate, a single genus (*Intestinibacter*) was associated in both cohorts. In contrast, associations with acetate and propionate were entirely cohort-specific. In order to control for delivery mode as a potential confounder, the analysis was re-run in a subset of vaginally born newborns (100% in the JP, and 57.1% in the PL, at birth). There were 25 genera associated with SCFA in the PL cohort (Figure 4C): acetate (N = 4), propionate (N = 4), butyrate (N = 10), and isobutyrate (N = 7). However the population-specific patterns persisted. For butyrate, two shared genera (*Intestinibacter* - associated in the same direction, and Incertae Sedis - mainly Ruminococcaceae - associated in the opposite direction) were identified. Associations with acetate, propionate and isobutyrate were entirely cohort-specific, with no overlapping genera identified (Figure 4D).These findings underscore the population-specific nature of microbial contributions to SCFA metabolism.

**Figure 4.**
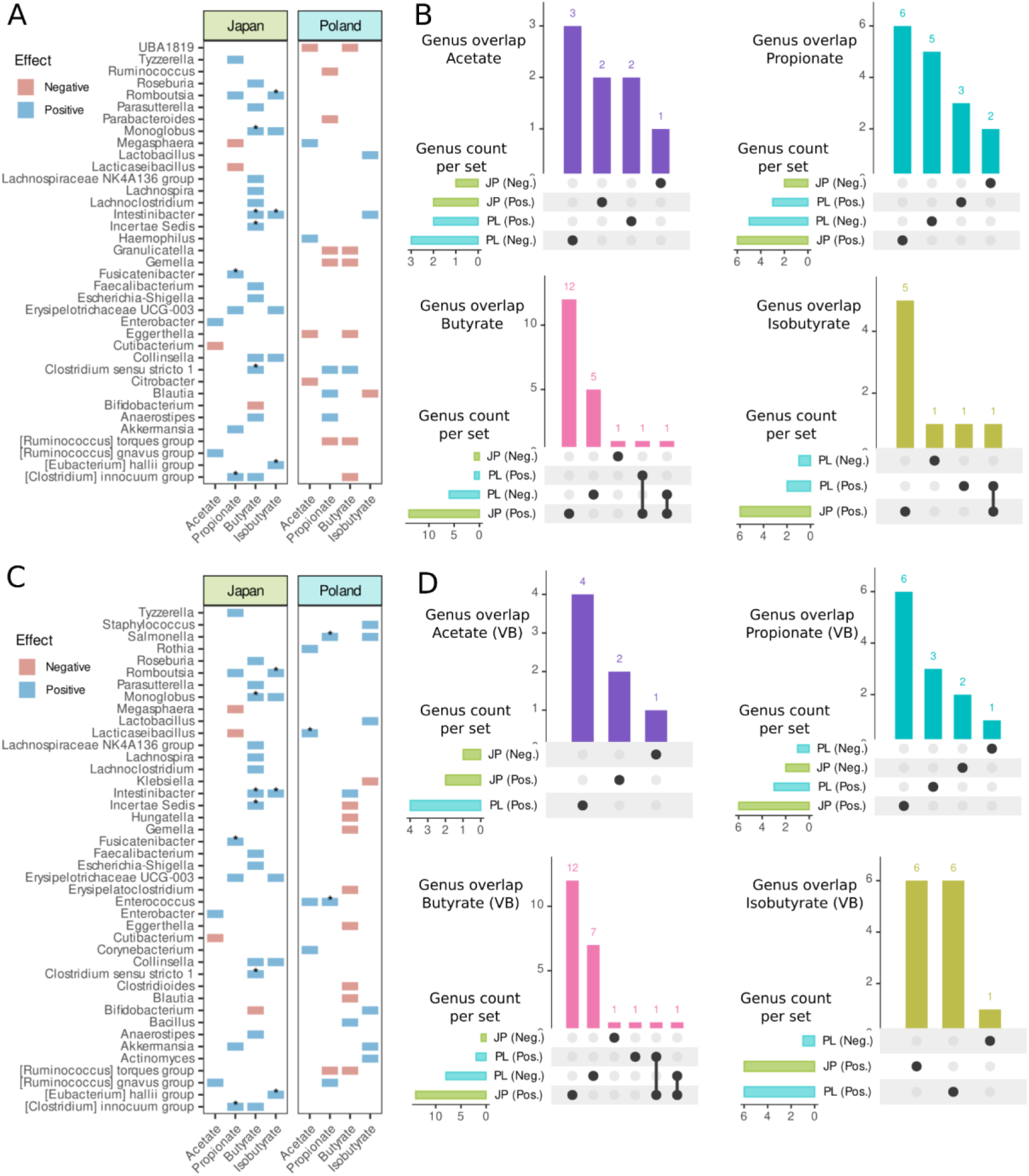
**Country-specific microbial associations with SCFAs and overlap of genus-level signals in full and vaginally-born cohorts** A - Direction of associations between microbial genera and concentrations of short-chain fatty acids (SCFAs) in Polish and Japanese cohorts. Each tile represents a genus–SCFA combination, colored by the direction of the effect (positive or negative). Only genera with significant associations (likelihood ratio test P < 0.05) are shown, and FDR adjusted significance is indicated with an asterisk. MIxed-effects models included the following covariates: sex, diet, antibiotic use and time point (as a categorical variable), SCFA was rank transformed; B - UpSet plot showing the overlap of genera significantly associated with each SCFA (P < 0.05), grouped by country (Poland vs. Japan) and effect direction (positive vs. negative). The intersection size bar plot indicates the number of genera shared across different country - effect combinations, while the set size horizontal bars represent the total number of genera per group: PL (Neg.) - Polish cohort, negative genus - SCF association, PL (Pos.) - Polish cohort, positive genus - SCFA association, LP (Neg.) - Japanese cohort, negative genus - SCF association, JP (Pos.) - Japanese cohort, positive genus - SCF association; C - Same as panel A, but restricted to the subset of vaginally-born (VB) newborns, to control for delivery mode as a confounding factor; D - Same as panel B, showing the overlap of significant genus - SCFA associations in vaginally-born (VB) newborns

To further investigate whether cohort-specific trajectories in genus-level abundance as well as distinct SCFA - genus associations translate into divergent metabolic outcomes, we next assessed SCFA dynamics over time by cohort. To unify SCFA concentrations reported in different units between the PL (nM/mg) and JP (mM) cohorts, we used z-score normalization which enabled comparability across cohorts while preserving within-cohort dynamics. The preservation of these dynamics can be observed in Figure 5A-D - raw SCFA trajectories in the PL (Figure 5A) and JP (Figure 5B) cohorts are maintained in their respective z-score normalized profiles (Figure 5C and Figure 5D). Interestingly, vaginally delivered newborns in the PL cohort exhibited more dynamic increases in acetate (sex, antibiotic use, and diet adjusted P value (P_adj_) = 0.045, Q = 0.112) and isobutyrate (P_adj_ = 0.056, Q = 0.112), as shown in Figure 5A. However, regardless of whether all or only vagianally delivered newborns were analysed, no significant differences in SCFA trajectories were observed between the PL and JP cohorts over the first 2 years of life. Model based predictions from mixed-effects linear models for all infants and for the vaginally delivered subset are shown in Figure 5E and Figure 5F, respectively. The obtained results suggest that despite observed differences in genus-level changes over time (Figure 3) and genus - SCFA associations (Figure 4) between the cohorts, SCFA profiles tend to remain relatively robust and preserved throughout the first two years of life.

**Figure 5.**
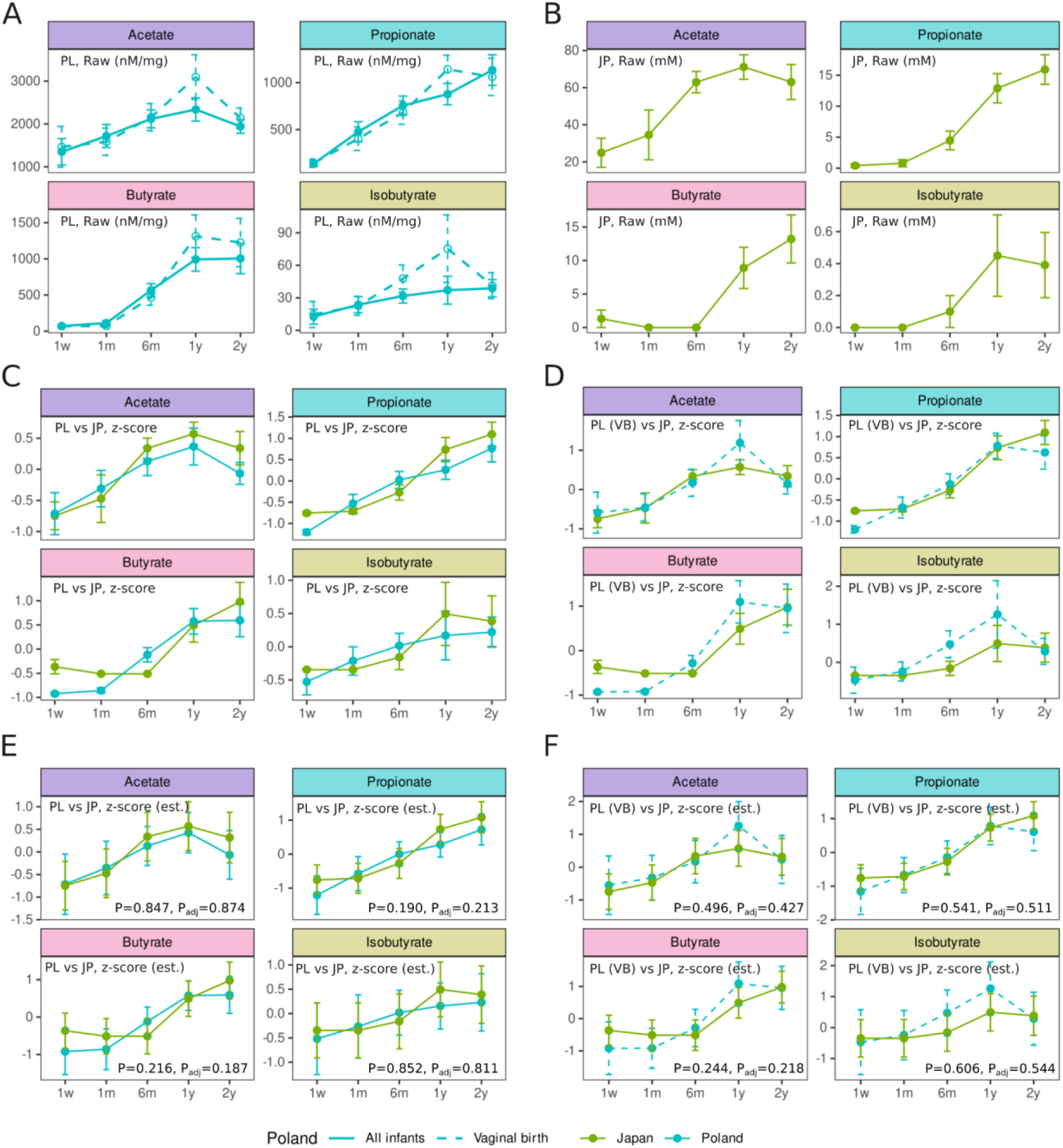
**SCFA dynamics over time in Polish and Japanese infant cohorts** A - SCFA concentrations over time in the Polish cohort (raw values), shown for all infants (solid line) and vaginally delivered (VB) infants (dashed line), B - SCFA concentrations over time in the Japanese cohort (raw values) - all Japanese infants were vaginally delivered, C - Z-score normalized SCFA trajectories in Polish vs. Japanese cohorts (all infants), D - Z-score normalized SCFA trajectories in Polish vs. Japanese cohorts (vaginally delivered only), E, F - Model-based predicted SCFA z-scores over time from linear mixed-effects models: all infants (E) and the vaginally delivered subset (F), P values for the time x cohort interaction obtained from likelihood ratio test comparing two nested models with and without the interaction term, P - Model without covariates, Padj - Model adjusted for sex, antibiotic use, and diet, PL - Polish cohort, JP - Japanese cohort, VB - vaginal birth, 1w - 1 week, 1m - 1 month, 6m - 6 months, 1y - 1 year, 2y - 2 years, est. - estimates - model-based predicted SCFA z-scores

To validate and extend our findings, we included two additional cohorts from diverse geographic regions - Finland (FIN) and the United States (US) - that met the inclusion criteria for gut microbiota comparisons (Supplementary Table 2). Although SCFA measurements were not available for these cohorts, limiting the ability to assess microbiota - metabolite (SCFA) associations, their inclusion enables both cross-sectional and longitudinal taxonomic analyses and strengthens the interpretation of findings obtained in the PL and JP cohorts. The ECAM project (US) includes data from six time points (N = 138 samples). The Finnish cohort includes data from five time points (excluding the 2-year time point, N = 516 samples). Metadata and time points for all four cohorts is available in Supplementary Table 2.

As a validation step, we extended the covariate-aware longitudinal analysis to the FIN and US cohorts to assess microbial trajectories over time. Country-specific changes in genus-level abundance, along with overlaps in significant genera identified using the full datasets, are shown in Supplementary Figures 13-14. Because the FIN and US cohorts included a larger number of participants and more complete sets of paired longitudinal samples than in the PL and JP cohorts, we additionally performed a subsampling analysis within each time interval to evaluate the robustness and reproducibility of temporal signals across balanced sample. The results of the subsampling analysis highlighting genera in the top 25% (upper quartile) of detection frequency (prevalence) across all genera are presented in Figure 6A. An UpSet plot, illustrating the directionality of genus-level abundance changes across consecutive and non-consecutive time intervals, is shown in Supplementary Figure 15. Only two genera exhibited discordant changes across cohorts: *Clostridium sensu stricto 1* (which increased in abundance from 6 months to 1 year in the JP cohort, but decreased in the FIN and US cohorts) and *Escherichia-Shigella* (which decreased in the JP cohort but increased in the FIN cohort between 1 week and 1 year; see Figure 6A). Given the low number of directionally inconsistent genera, Figure 6B presents Venn diagrams that visualize intersections of all significant genus-level changes between consecutive time points across cohorts, irrespective of the direction of change (for all consecutive and non consecutive intervals see Supplementary Figure 16).

**Figure 6.**
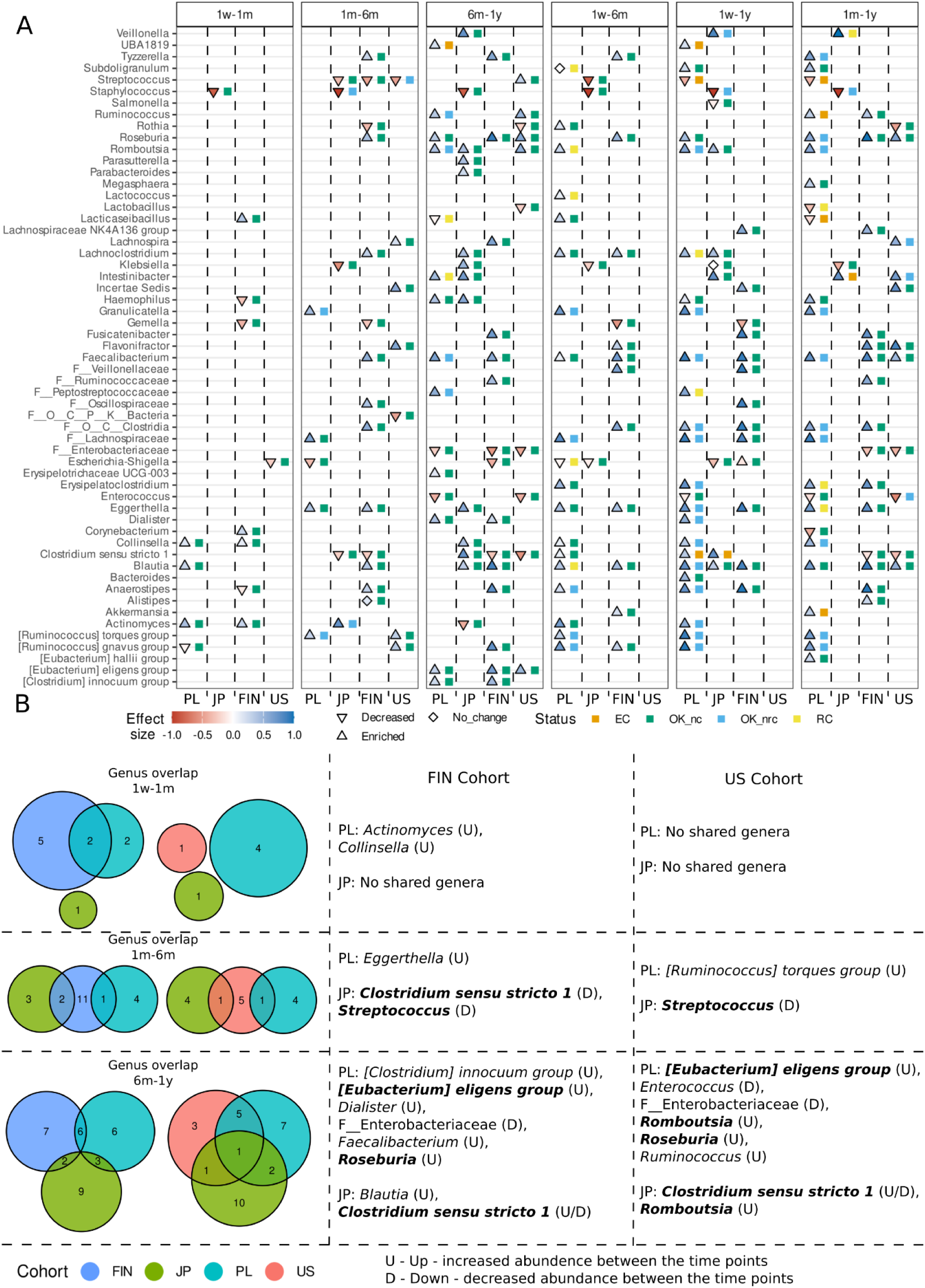
**Cohort-specific significant changes in genus-level abundance across time points (consecutive and non-consecutive) in PL and JP (full cohorts) and in subsampled FIN and US cohorts**

## Discussion

This study provides a longitudinal analysis of the infant gut microbiota across 4 cohorts (PL, JP, FIN, US), showing mostly cohort-specific temporal microbial signatures during the first two years of life (Figure 3). Despite marked between-cohort differences in microbiome taxonomy our validation analyses identified some genera consistently shared across cohorts, including *Faecalibacterium*, *Blautia*, *Anaerostipes*, *Eggerthella*, *Ruminococcus*, and members of the *Lachnospiraceae* family. The appearance of these taxa aligns with the “core” age-predictive species identified by Bottino et al. (2024), where *Faecalibacterium prausnitzii*, *Anaerostipes hadrus*, *Ruminococcus gnavus* and *Blautia wexlerae* emerged as the important predictors of microbiome maturation across 12 countries. The degree of overlap in our study was especially evident between the PL and FIN or US cohorts, whereas the JP cohort showed fewer shared taxa (Figure 6, Supplementary Figure 16).

The present study also provides novel insights into the temporal dynamics of SCFA production in early life, a critical window for microbiota maturation and host development (Lynch et al., 2023). A central and novel finding of our work is the demonstration that children in geographically distinct cohorts (Japanese and Polish) exhibited remarkably similar SCFA profiles despite substantial differences in both taxonomic composition (Figure 3) and taxa- SCFA associations (Figure 4). To our knowledge, this represents the first *in vivo* evidence that SCFA production during infancy follows a highly conserved developmental trajectory across geographically and culturally distinct populations. This observation strongly supports the concept that the functional maturation of the infant gut microbiome is more tightly constrained than its taxonomic composition, underscoring the notion that distinct microbial communities can converge functionally to ensure consistent metabolic outputs during this key developmental period.

### Functional Convergence Despite Taxonomic Divergence in Early-Life SCFA Metabolism

To date, only a limited number of studies have comprehensively investigated SCFA production during early life (Midtvedt and Midtvedt, 1992; Norin et al., 2004; Differding et al., 2020; Łoniewska et al., 2023; Nilsen et al., 2024). Nilsen et al. (2024) examined the temporal development of infant gut microbiota and fecal SCFA profiles from meconium to 12 months in relation to immune cell composition, but did not include direct analyses linking microbial taxa to SCFA levels. In contrast, Norin et al. (2004) compared fecal SCFA concentrations in Estonian and Swedish infants and found that Swedish infants had significantly higher SCFA levels at one month, nevertheless both populations displayed a similar shift toward higher-chain fatty acids (C4-C6) by 12 months. Differding et al. (2020), while incorporating 16S rRNA sequencing, only conducted cross-sectional correlation analyses between taxa and SCFAs at two discrete time points (3 and 12 months) and measured only relative SCFA abundances. In this study, gut microbiota and SCFA were analysed longitudinally over 24 months at 5 time points, and taxa-SCFA associations were examined using linear mixed-effects models. Although both cohorts showed numerous statistically significant associations between genera and SCFA concentrations (Figure 4), the overlap was minimal. For instance, only *Clostridium sensu stricto 1* and the *[Clostridium] innocuum* group were associated with butyrate in both cohorts, and *Intestinibacter* was the only genus consistently linked to isobutyrate. Notably, taxa associated with acetate and propionate were entirely cohort-specific, suggesting population-level divergence in microbial contributors to SCFA metabolism. Strikingly, however, after z-score normalization of SCFA concentrations, trajectories for acetate, propionate, and butyrate were nearly identical between cohorts and remained stable after adjustment for early-life confounders (Figure 5). Although early-life exposures such as delivery mode, perinatal antibiotics, and feeding type modulate microbiota composition and in consequence SCFA levels (Midtvedt and Midtvedt, 1992; Łoniewski et al., 2022b; Bui et al., 2025), they appear to exert limited influence on the overall progression of SCFA production.

Our observations strongly support the concept of functional redundancy, in which different microbial taxa fulfill overlapping metabolic roles. Functional redundancy - defined as the capacity of distinct species to contribute similarly to ecological functions - is increasingly recognized as a key feature of healthy microbiomes along with resistance resilience and stability (Moya and Ferrer, 2016). In early life, Nguyen et al. (2021) reported that taxonomic profiles poorly predicted stool metabolomic signatures, suggesting the buffered and bidirectional nature of microbiome - metabolome relationships. Cerdó et al. (2018) further demonstrated that SCFA-associated proteins remained stable over time despite shifting taxonomic compositions, while Guittar et al. (2019) showed that trait-based microbial signatures stabilized earlier than OTU-level taxonomic profiles, reinforcing the notion that function can persist despite taxonomic turnover. Mechanistic evidence also supports this paradigm. Reichardt et al. (2018) showed that in vitro fermentation of 15 non-digestible carbohydrates (NDCs) by fecal microbiota from different individuals produced remarkably consistent SCFA profiles for each substrate, despite large interindividual differences in microbial composition. However, they also noted that metabolites produced by a narrow range of taxa were more variable, and that the absence of key species - such as *Ruminococcus bromii* in resistant starch fermentation - significantly diminished SCFA output. This suggests that functional buffering has limits and may depend on the presence of keystone taxa. Further support comes from Appert et al. (2020), who investigated the establishment of butyrate-producing bacteria in 16 Swiss infants sampled at 1, 3, and 12 months of age. They demonstrated that early butyrate production was seeded predominantly by endospore-forming *Clostridium sensu stricto*, which are metabolically less efficient, and later complemented by high-efficiency butyrate producers such as *Faecalibacterium prausnitzii*,

*Roseburia/Eubacterium rectale*, and *Anaerobutyricum hallii*. Importantly, they suggested that coexisting Clostridiaceae, Ruminococcaceae and Lachnospiraceae ensure functional redundancy within a given infant gut community. While these findings support the ecological principle that conserved functional outputs can emerge from divergent microbial communities, it is important to note that Appert’s sampling design included three time points (1, 3, and 12 months), only two of which overlapped with our PL and JP cohorts. For this reason, their dataset was not included in our reanalysis, and a direct comparison of SCFA trajectories with our cohorts was not feasible. Recently, Fahur Bottino et al. (2025) identified a small group of taxa - including *Faecalibacterium prausnitzii* and *Anaerostipes hadrus* - that consistently appear in succession across diverse geographic and socioeconomic settings and serve as robust predictors of infant age (“core” age-predictive species). In our data, both taxa were positively associated with butyrate, but only in the Japanese cohort, while *Bifidobacterium* was negatively associated. This population-specific variation in taxon - function associations suggests that, although some microbes may be globally conserved, their metabolic roles are context-dependent and shaped by factors such as diet, host genetics, and environmental exposures. Taken together, our findings highlight a fundamental principle: while taxonomic profiles of the infant gut microbiome diverge across populations, metabolic outputs essential for host development - such as SCFA production - follow a conserved and resilient trajectory.

### Cross-sectional biodiversity and taxonomy across early developmental stages

Our cross-sectional analyses revealed notable differences in microbial diversity between Japanese and Polish newborn cohorts. While most alpha-diversity indices (evenness, Shannon, Simpson) showed no statistically significant differences, trends consistently indicated higher microbial diversity in the Polish cohort, particularly evident at six months of age. Richness appeared as the most sensitive index, with a significant difference at one week and a reproducible divergence across multiple time points when analyses were restricted to ASVs mapped to taxa shared between both cohorts. When restricting analyses to the 74 genera shared between cohorts, overall alpha diversity was significantly reduced compared to unrestricted metrics, suggesting that cohort-specific or unassigned taxa inflated apparent diversity estimates. The restricted analysis, however, revealed consistent and statistically significant differences between the cohorts across nearly all time points, suggesting that both unique and shared taxa contribute to shaping distinct microbial ecologies in Polish and Japanese infants. Importantly, resampling approaches confirmed that these observations were not artifacts of unequal sample sizes or sequencing depth, reinforcing the robustness of these cohort-level distinctions. These findings are consistent with previous longitudinal cohort studies showing that microbial diversity steadily increases during the first years of life but remains lower than in adults until well beyond infancy. In a large Swedish cohort followed up to five years, Roswall et al. (2021) demonstrated that alpha diversity rises continuously with age, though even at age five, diversity had not yet reached adult levels. Similarly, Tsuruoka et al. (2024) confirmed significant stepwise increases in Shannon diversity between 4, 10, 18 months, and 3.5 years, accompanied by gradual convergence toward an adult-like microbial configuration. Together with our data, these studies suggest that geographic and cultural factors may shape early richness, yet the overall trajectory of diversity accumulation is a conserved feature of early microbial ecology.

Beta-diversity analyses further highlighted consistent and significant country-specific differences in microbial community structure, persisting after adjustment for major covariates such as sex, diet, and antibiotic exposure. These findings were largely reproduced when analyses were restricted to shared genera, indicating that differences in community composition were not solely driven by cohort-specific taxa, but rather by divergent structuring of a shared microbial core. This is in line with recent analyses of global cohorts, which have demonstrated that while the identity of key colonizing taxa is conserved, their relative abundances and timings of succession can differ between populations (Fahur Bottino et al., 2025). While no single genus was consistently enriched across all ages, several patterns emerged. The majority of differentially abundant genera were enriched in the Polish cohort, suggesting a broader taxonomic expansion relative to Japanese infants. Conversely, a subset of genera, including *Staphylococcus* in early infancy, *Klebsiella* at one month, and *Bifidobacterium* and *Enterococcus* at later stages, were significantly more abundant in the Japanese cohort. Interestingly, features not assigned to a defined genus but classified at higher taxonomic levels (e.g., *Bifidobacteriaceae*) were consistently enriched in the Polish infants, highlighting the potential influence of early-life bifidobacterial dynamics. This observation is particularly relevant given the well-established role of bifidobacteria in metabolizing human milk oligosaccharides (HMOs) and producing acetate as a cross-feeding substrate for butyrate-producing taxa (Marcobal et al., 2011; Rivière et al., 2016; Nguyen et al., 2021). Notably, the competitive or cooperative interactions between bifidobacteria and butyrate producers may differ depending on strain composition, which could underlie some of the cohort-specific divergences in microbial structure.

Sensitivity analyses restricted to vaginally delivered infants confirmed the robustness of the primary findings. Alpha- and beta-diversity differences, as well as the overall structure of differential abundance results, were consistent with those observed in the full cohorts. However, some genera that were significantly different in the broader analysis lost significance in the vaginally born subset, while a previously unreported taxon group (*Burkholderia– Caballeronia–Paraburkholderia*) emerged as differentially abundant. These observations suggest that delivery mode, while not independently comparable across cohorts due to the lack of cesarean-born infants in Japan, may modulate specific taxonomic signals within the broader cohort-level differences. Prior studies have also shown that delivery mode has large effects on microbial composition early in life, with cesarean-born infants exhibiting reduced alpha diversity and delayed acquisition of certain taxa, though these differences diminish by later childhood (Dominguez-Bello et al., 2010; Roswall et al., 2021) . Taken together, these findings demonstrate that Polish and Japanese infants may harbor distinct microbial ecosystems early in life, shaped both by shared microbial cores and population-specific expansions.

### Cohort-specific genus trajectories dominate early-life microbiome maturation

Longitudinal analyses revealed striking differences in microbial trajectories between the Polish and Japanese infant cohorts. Although both groups exhibited significant within-cohort temporal changes, only a small subset of genera displayed consistent trajectories across populations. For example, *Staphylococcus*, *Romboutsia*, *Intestinibacter*, and *Haemophilus* were shared between the cohorts in specific intervals, yet the majority of genus-level dynamics were unique to each population. This pattern persisted even when restricting analyses to vaginally born Polish infants, confirming that delivery mode alone does not account for these divergent developmental patterns. Genera with shared longitudinal shifts were relatively rare, accounting for less than 20% of all significantly altered taxa in either cohort, underscoring the predominance of country-specific ecological trajectories during the first two years of life. While additional genera such as *Lachnoclostridium*, *Clostridium sensu stricto 1*, *Lactococcus*, and *Flavonifractor* were occasionally identified as shared when longer intervals were considered, these represented exceptions rather than the rule. Together, these findings emphasize that while a small number of core taxa appear to follow conserved temporal dynamics across populations, the majority of microbial shifts during early development are contingent on cohort-specific factors, likely reflecting environmental, cultural, and dietary influences. These results align with ecological succession models of the infant microbiome, in which early colonization by taxa such as *Bifidobacterium* and *Enterobacteriaceae* is gradually supplanted by butyrate producers like *Faecalibacterium* and members of the *Lachnospiraceae* family (Roswall et al., 2021; Fahur Bottino et al., 2025). Such succession processes, while broadly conserved, may progress along slightly different temporal trajectories across populations, contributing to the cohort-specific signals observed in our analyses.

This interpretation is supported by evidence from European cohorts, where early and diverse complementary feeding accelerates microbial maturation. Dikareva et al. (2025), studying Dutch infants, showed that early introduction of fruits and vegetables was associated with higher microbial diversity at four months, earlier enrichment of butyrate producers such as *Flavonifractor plautii*, and smoother transitions in community structure during the weaning window. These infants also demonstrated greater functional potential for fiber degradation and butyrate synthesis, consistent with accelerated ecological succession. Similarly, Roswall et al. (2021) found that the infant microbiome follows distinct trajectories during the first five years, with bifidobacteria dominating in early infancy, followed by the gradual expansion of *Bacteroides*, *Faecalibacterium*, and *Roseburia* after the introduction of solid foods. Tsuruoka et al. (2024) further demonstrated that microbiota diversity increases stepwise from 4 months to 3.5 years, accompanied by compositional convergence toward the maternal microbiome. Taken together, these findings indicate that while the broad succession pattern is conserved, differences in diet timing, diversity, and cultural practices shape the pace of maturation.

### Strengths and limitations

The strengths of our study lie in its robust methodological design, characterized by longitudinal sampling spanning two critical years of infant development, standardized protocols, stringent statistical control of confounding variables, and the integration of microbiota composition with SCFA metabolite profiles. Additionally, our study leveraged publicly available datasets for validation across cohorts, enhancing the generalizability and robustness of the findings.

Despite the comprehensive nature of our analysis, our study has several important limitations. First, the relatively modest and imbalanced sample size, particularly at later time points such as two years, limited the statistical power of our analyses and may have affected the generalizability of our conclusions. The selection of samples at predefined intervals and the loss of some participants over time further exacerbated this issue, potentially introducing bias into the longitudinal assessment. To address the challenge of modest and imbalanced sample sizes, particularly at later time points, we harmonized sampling intervals across cohorts by selecting time-point-matched samples with a clearly defined and consistent tolerance range (±10% age tolerance) when exact matches were unavailable.

Second, the nature of our study as a post hoc analysis, involving the combining and comparing of separately collected datasets, rather than conducting a prospective, comparative longitudinal study. This methodological approach carries a significant risk of bias arising from differences in recruitment strategies, inclusion and exclusion criteria, clinical management practices, laboratory protocols, and data collection standards across cohorts. Such discrepancies may limit the accuracy and interpretability of comparative analyses. Recognizing that our study was a post hoc comparative analysis utilizing datasets collected independently, we implemented validation strategies using two additional cohorts (FIN and US cohorts) to confirm methodological consistency and analytical reproducibility. This validation step partially mitigated biases arising from differences in protocols, and clinical management. However, this validation cannot completely replace the methodological rigor provided by a prospective comparative study design. Therefore, future research should prioritize prospective, multicenter comparative longitudinal studies using standardized inclusion and exclusion criteria, sampling schedules, clinical and laboratory protocols.

Third, our analysis was based on 16S rRNA gene amplicon sequencing, a widely used but inherently limited method, which restricts resolution to genus-level taxonomy. In addition, variability in targeted hypervariable regions of the 16S rRNA gene across the included cohorts represents a notable methodological constraint. These differences could introduce biases in diversity estimates, and compositional comparisons, potentially influencing the accuracy of cross-cohort comparisons and limiting direct taxonomic and functional inferences. To reduce biases introduced by variability in targeted hypervariable regions (V1-V2, V3-V4, and V4), we applied a uniform bioinformatic workflow across cohorts, including a standardized rarefaction depth and harmonized preprocessing and filtering protocols. Specifically, we used identical analytical parameters for diversity calculations (richness, evenness, Simpson index, and Shannon index) and performed additional analyses restricted to amplicon sequence variants (ASVs) mapped to genera common between the main cohorts (Polish and Japanese). This approach addresses potential biases in compositional comparisons.

Fourth, the public cohorts (JP, FIN, and US) exhibited inherent heterogeneity in terms of sample processing protocols, sampling frequencies, sequencing platforms, and metadata quality. We addressed the inherent heterogeneity across public cohorts by rigorously harmonizing metadata, sampling intervals, and microbiome data prior to statistical analyses. Data harmonization included uniform age binning, selection criteria for samples at each time point, standardized preprocessing steps, and rigorous quality control. The identification of consistent biological signals across cohorts, despite notable differences in methodology, technology, and data quality, strengthens the credibility of our findings and supports the interpretation that the observed effects are biologically meaningful rather than technical artifacts.

Finally, our SCFA analyses were constrained by methodological differences and availability between cohorts. Specifically, SCFA measurements were only available in the Polish and Japanese cohorts and were reported using distinct measurement units. We mitigated these discrepancies by z-score normalization. While z-score normalization facilitates comparative analyses, it obscures absolute differences in metabolite levels, potentially masking biologically relevant variations in metabolite concentrations across populations. Importantly, we used z-score normalization only for comparative analyses between cohorts, Polish and Japanese, which enabled meaningful statistical analyses. However, for intra-cohort analyses examining temporal dynamics, we deliberately retained the original SCFA measurement units to preserve absolute quantification and enable precise interpretation of within-cohort metabolic changes.

## Conclusions

This is the first comprehensive re-analysis study to characterize and directly compare the development of gut microbiota and fecal short-chain fatty acid (SCFA) profiles in healthy Polish and Japanese infants during the first two years of life. Our findings highlight a nuanced pattern in the associations between bacterial genera and SCFA concentrations, revealing that some putative SCFA producers are shared across populations whereas others are population-specific. Most notably, and constituting the principal novelty of this work, both cohorts exhibited the same age-related pattern of change in stool SCFA concentrations. Despite marked, cohort-specific taxonomic trajectories, model-based trajectories of acetate, propionate, butyrate and isobutyrate were conserved between populations, indicating convergent functional maturation of the gut ecosystem. To our knowledge, such cross-population convergence of SCFA dynamics in early life has not been previously reported.

These results demonstrate fundamental principle of infant gut ecology: while taxonomic profiles and genus-SCFA relationships diverge, core metabolic functions crucial for host development follow a conserved and resilient trajectory. This has important implications for early-life microbiome research and interventions, suggesting that supporting functional maturation may be more relevant for host health than targeting specific taxa. Future work integrating host growth, immune, and neurodevelopmental outcomes with microbial functional trajectories, and that include a wider representation of cohorts, will help fully understand the role of conserved SCFA dynamics in shaping long-term health.

### Data availability

The datasets used and/or analysed during the current study are available on reasonable request ECAM: PRJEB14529, Japanese: PRJDB9469, Finnish: PRJEB48451

## Supporting information

Supplemental Figures & Tables descriptions

