## Supplemental Figures & Tables descriptions for "Similar Fecal SCFA Patterns Despite Diverse Gut Microbiota in Polish and Japanese Children During the First Two Years of Life. The longitudinal, comparative, validated study"

**Supplementary Figures**

**Supplementary Figure 1.** SCFA levels over time (with age bin ranges) in JP cohort
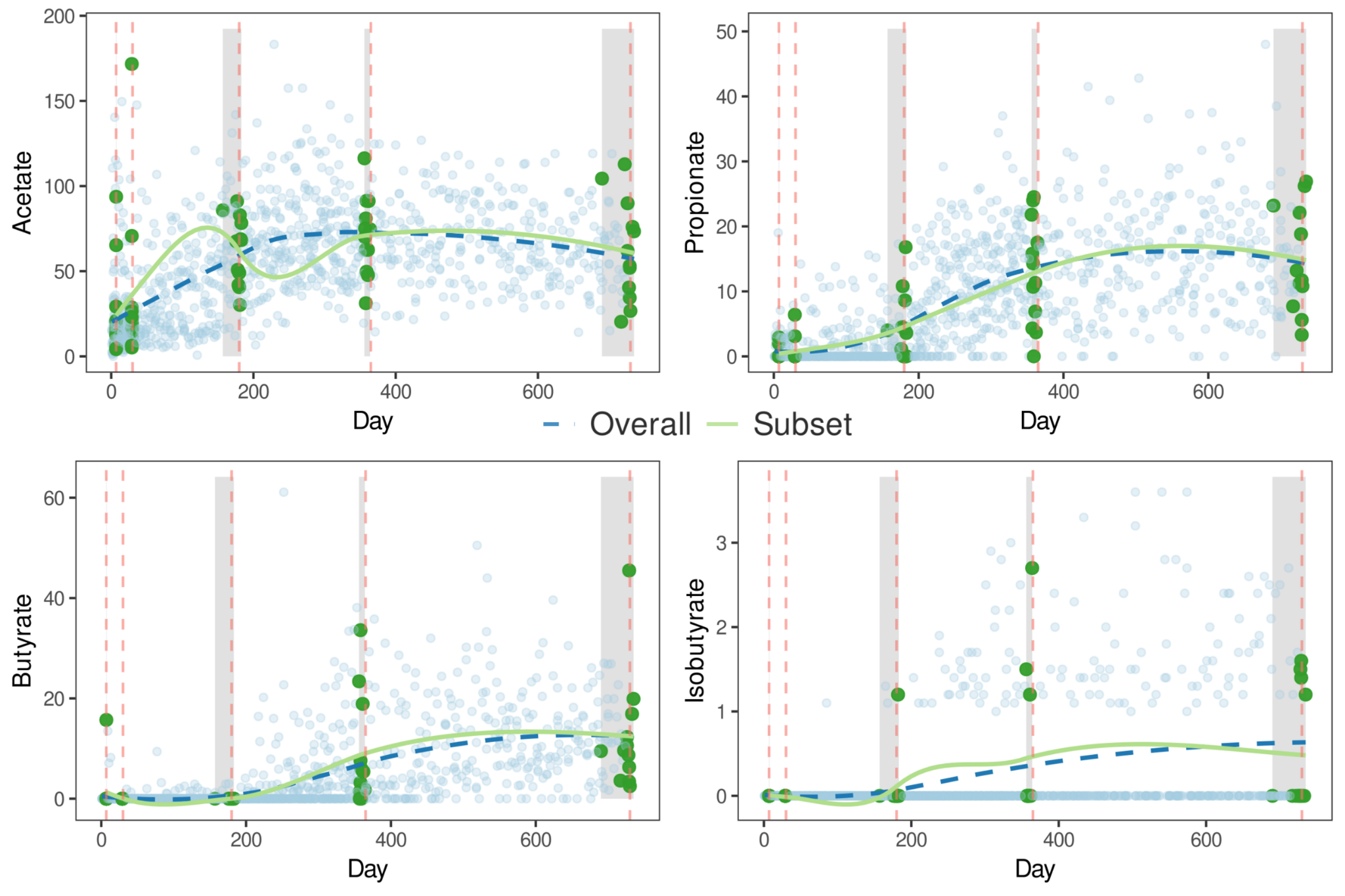

The subset of JP samples selected for this study is indicated in green, while blue points represent all 1,048 samples (see Supplementary Table 3) from the longitudinal study of 12 children by Tsukuda et al. (Tsukuda et al., 2021). Light gray zones indicate the time ranges corresponding to each age bin, and vertical red dashed lines denote approximate standard age mappings (7, 30, 180, 365, and 730 days). Smoothed trends were derived using LOESS fitting: the blue dashed line reflects the overall trend (all samples), while the green solid line corresponds to the selected subset. Age bin ranges for JP samples (in days) were: 7–7.5 (1st week), 29 (1 month), 157–183 (6 months), 356–364 (1st year), and 690–735 (2nd year). In the PL cohort, bins included the first stool and the 1st week (7 days, collected at the hospital), 30 and 180 days ±3 days (1st and 6th month), and 365 and 730 days ±7 days (1st and 2nd year).

**Supplementary Figure 2**. Alpha-diversity at each time point and by country (Japan *versus* Poland) - non-restricted

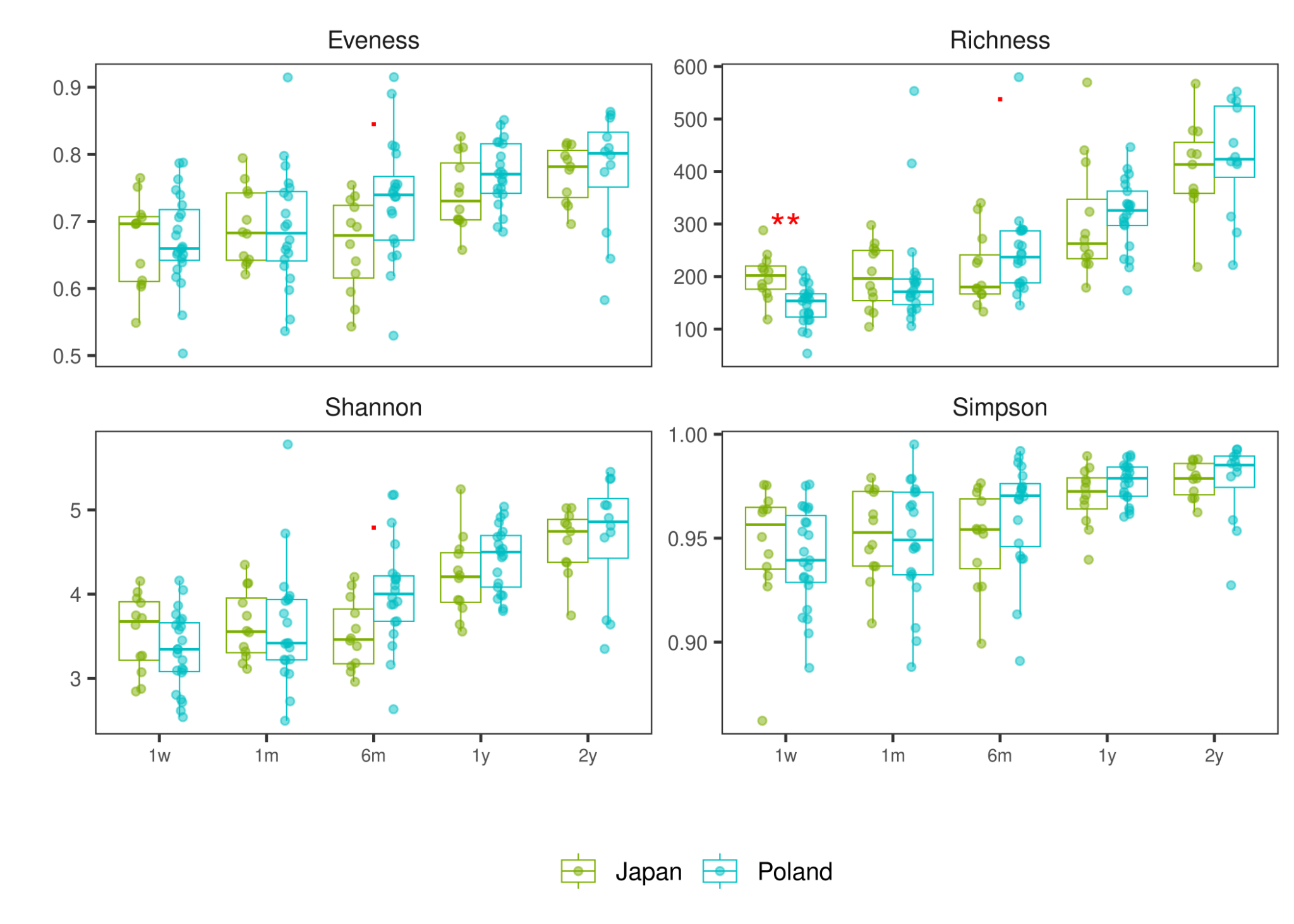
Cross-sectional comparisons between the PL and JP cohorts were conducted using linear models that included antibiotic exposure (ATB), sex, and diet as covariates, except at the 2-year timepoint, where none of the newborns were breastfed. P values were generated using ANOVA comparing two nested models - one including the country term and covariates, and one excluding the country term. Multiple testing was controlled using the false discovery rate (FDR) procedure applied separately at each time point. Boxplots and points (raw data) display the distribution of alpha diversity metrics by country and time point.

Statistical significance is denoted by stars according to FDR-adjusted P values: 0 - 0.001 '***', 0.001 - 0.01 '**', 0.01 - 0.05 '*', 0.05 - 0.1 '.', 0.1 - 1.0 (No symbol)

1w - 1 week of life, 1m - 1 month, 6m - 6 months, 1y - 1 year, 2y - 2 years.

**Supplementary Figure 3**. ASVs statistics on resampling to equal sample size (25 - 50) over different number of iterations

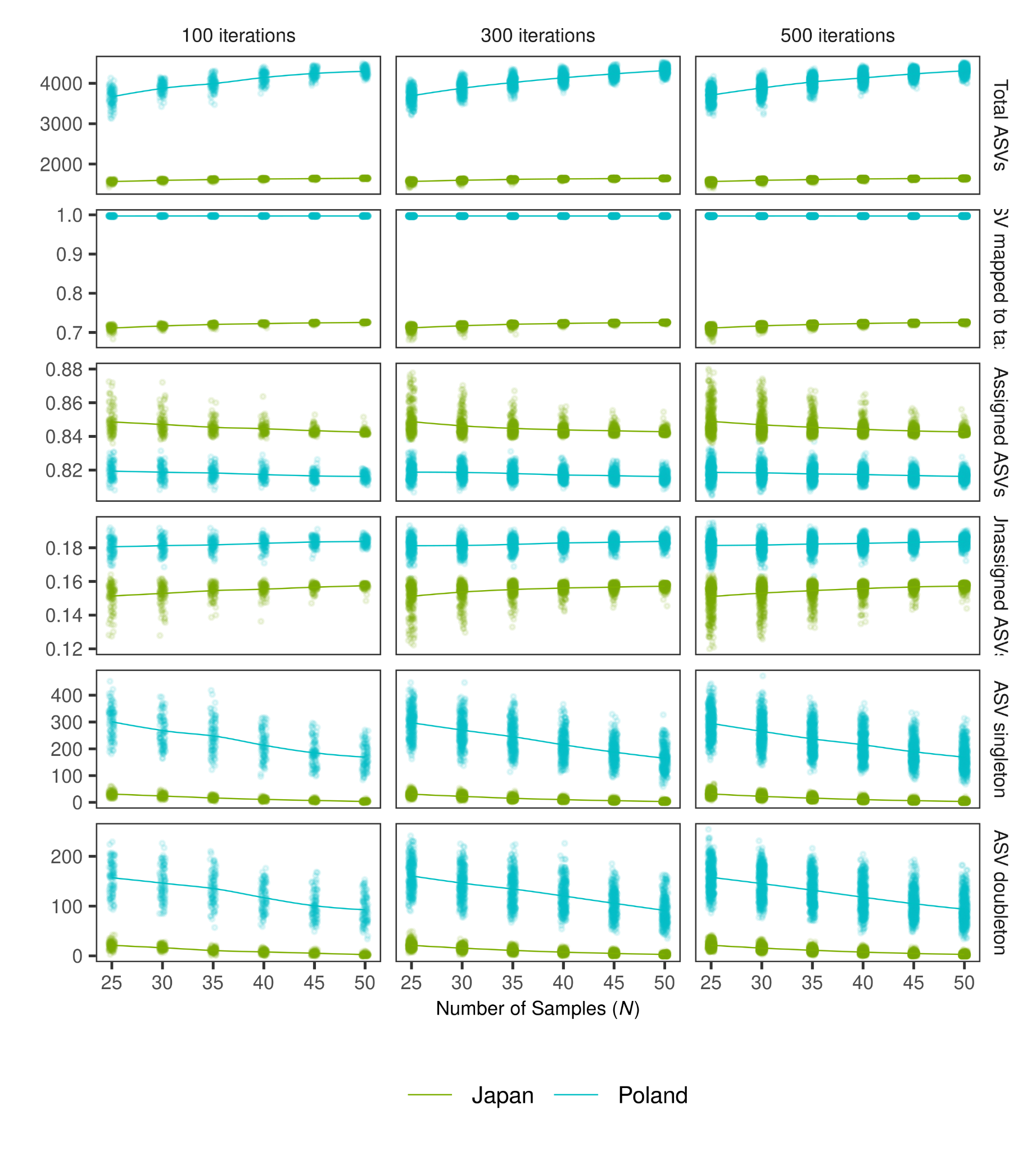

Diversity and mapping statistics of ASVs (Amplicon Sequence Variants) in random and equal subsets of microbiome data across two cohorts (Poland and Japan). For each random subset the following statistics were assessed: total ASVs, ASVs mapped to taxonomy at the genus level, and proportions of assigned/unassigned taxa at the genus level, singletons (ASVs detected only in one of the selected samples) and doubletons (ASVs with a total count of 2 across selected samples). Repeated for various values of sample number (N) and number of resamplings (iterations). As the number of individuals (N) included in each resample increased, the number of singletons and doubletons (ASVs observed only once or twice) gradually decreased. This trend is expected - when more samples are pooled together, the likelihood of an ASV being seen only once (or twice) decreases. Other metrics - total number of observed ASVs, proportion of ASVs mapped to the taxonomic hierarchy, proportion of assigned vs. unassigned taxa remained relatively stable across the range of tested sample sizes. The differences between the Poland and Japan cohorts in terms of ASV richness, mapping rates, and assignment rates were relatively consistent across sample sizes, thereby implying that cohort-level differences are not driven by sampling depth or sample size.

**Supplementary Figure 4**. Alpha-diversity at each time point and by country (Japan *versus* Poland) - the analysis restricted to ASVs mapped to a set of 74 genera shared between cohorts

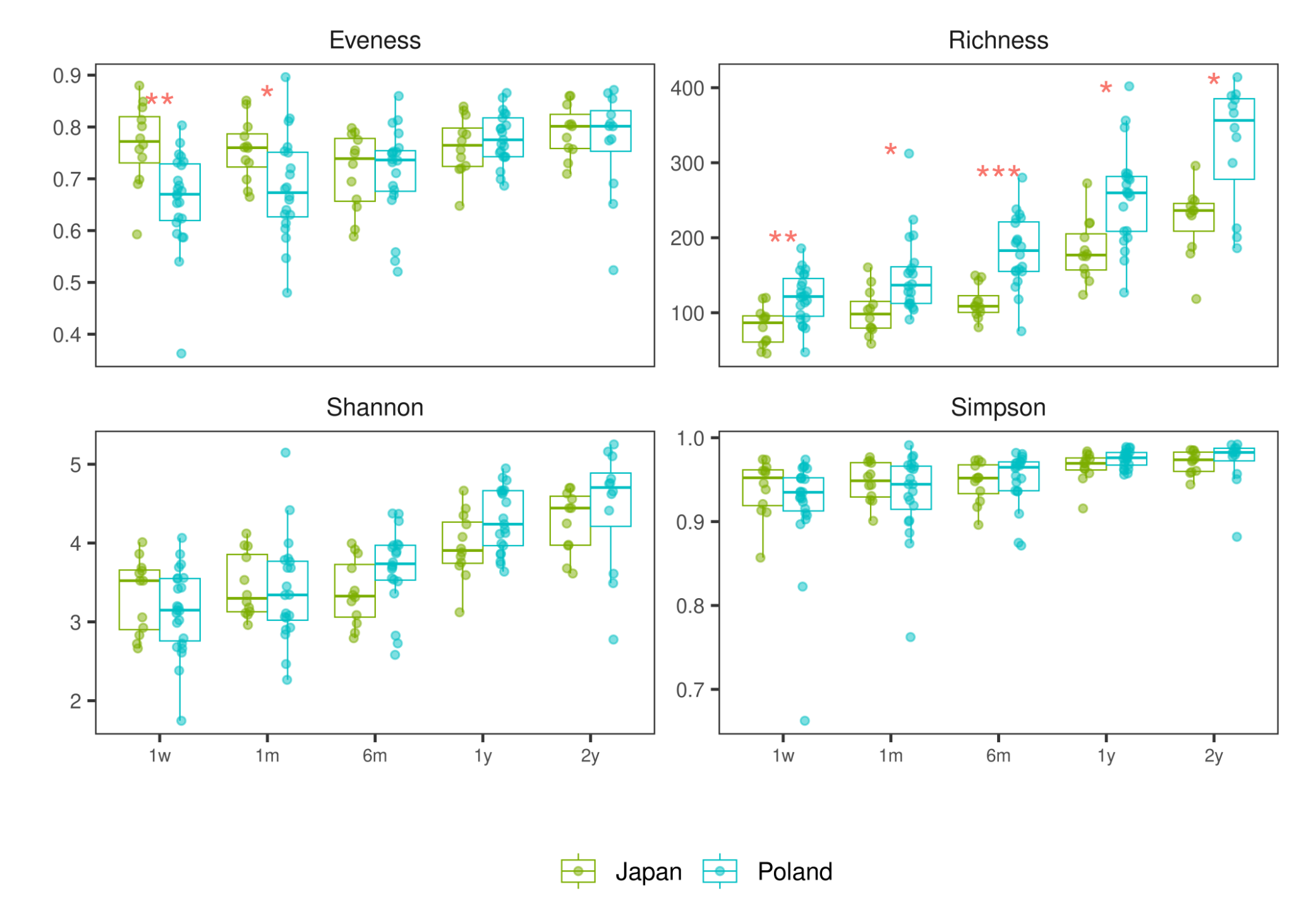

Cross-sectional comparisons between the PL and JP cohorts were conducted using linear models that included antibiotic exposure (ATB), sex, and diet as covariates, except at the 2-year timepoint, where none of the newborns were breastfed. P values were generated using ANOVA comparing two nested models - one including the country term and covariates, and one excluding the country term. Multiple testing was controlled using the false discovery rate (FDR) procedure applied separately at each time point. Boxplots and points (raw data) display the distribution of alpha diversity metrics by country and time point.

Statistical significance is denoted by stars according to FDR-adjusted P values: 0 - 0.001 '***', 0.001 - 0.01 '**', 0.01 - 0.05 '*', 0.05 - 0.1 '.', 0.1 - 1.0 (No symbol)

1w - 1 week of life, 1m - 1 month, 6m - 6 months, 1y - 1 year, 2y - 2 years.

**Supplementary Figure 5**. Alpha-diversity at each time point and by country (Japan *versus* Poland) - **only vagianally born**, non-restricted

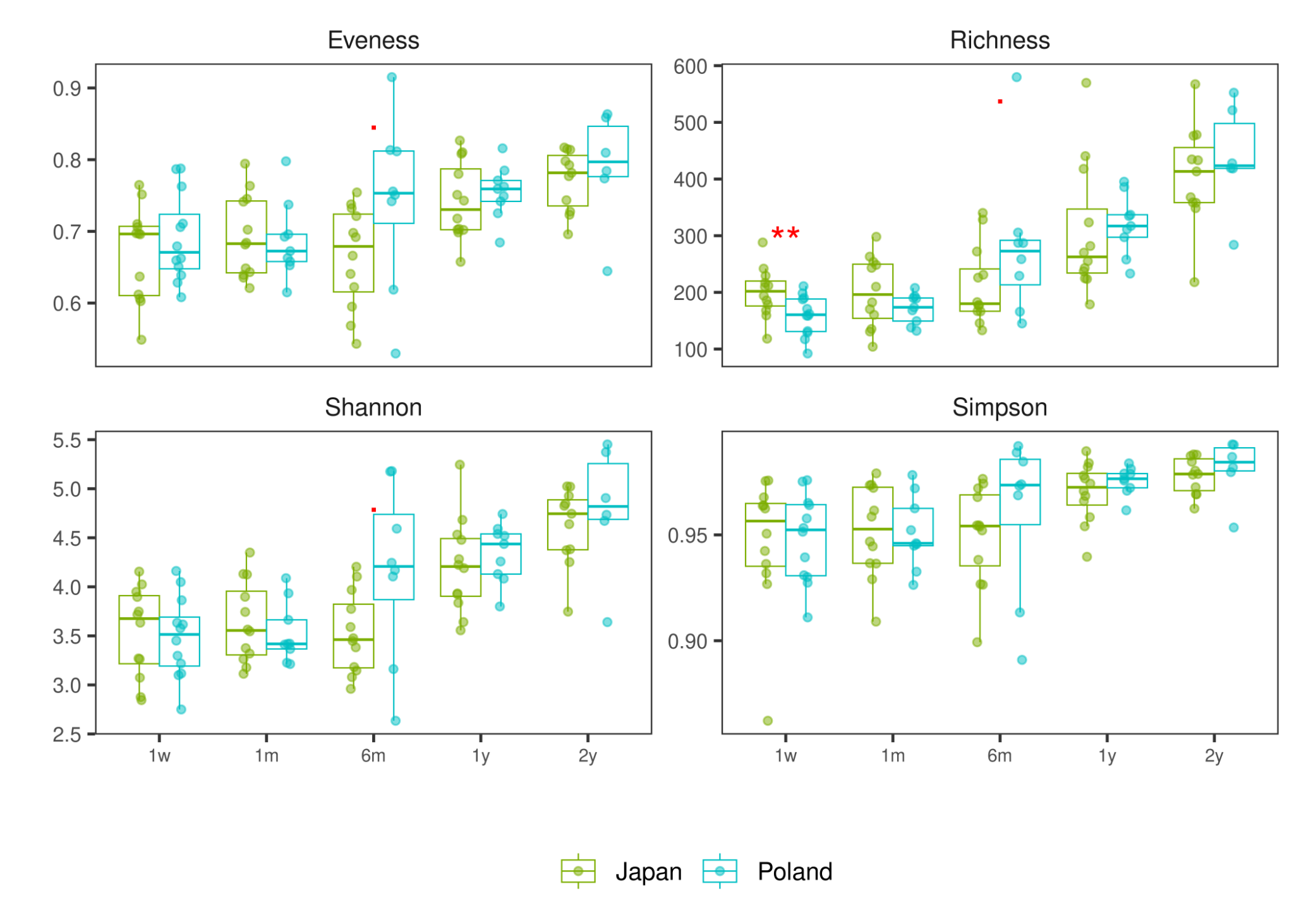
Cross-sectional comparisons between the PL and JP cohorts were conducted using linear models that included antibiotic exposure (ATB), sex, and diet as covariates, except at the 2-year timepoint, where none of the newborns were breastfed. P values were generated using ANOVA comparing two nested models - one including the country term and covariates, and one excluding the country term. Multiple testing was controlled using the false discovery rate (FDR) procedure applied separately at each time point. Boxplots and points (raw data) display the distribution of alpha diversity metrics by country and time point.

Statistical significance is denoted by stars according to FDR-adjusted P values: 0 - 0.001 '***', 0.001 - 0.01 '**', 0.01 - 0.05 '*', 0.05 - 0.1 '.', 0.1 - 1.0 (No symbol)

1w - 1 week of life, 1m - 1 month, 6m - 6 months, 1y - 1 year, 2y - 2 years.

**Supplementary Figure 6**. Alpha-diversity at each time point and by country (Japan *versus* Poland) - the analysis restricted to ASVs mapped to a set of 74 genera shared between cohorts, **vaginally-born newborns**

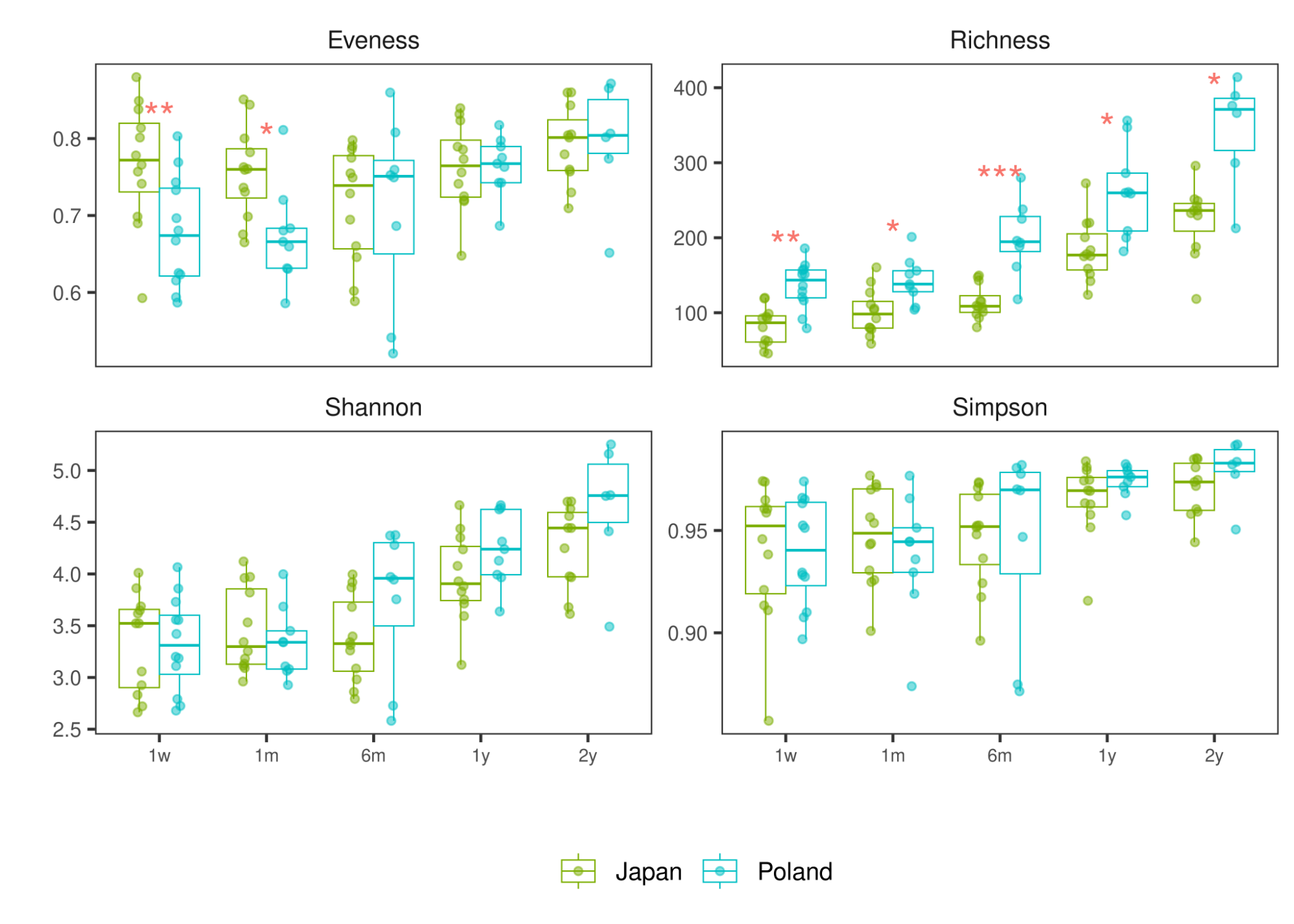

Cross-sectional comparisons between the PL and JP cohorts were conducted using linear models that included antibiotic exposure (ATB), sex, and diet as covariates, except at the 2-year timepoint, where none of the newborns were breastfed. P values were generated using ANOVA comparing two nested models - one including the country term and covariates, and one excluding the country term. Multiple testing was controlled using the false discovery rate (FDR) procedure applied separately at each time point. Boxplots and points (raw data) display the distribution of alpha diversity metrics by country and time point.

Statistical significance is denoted by stars according to FDR-adjusted P values: 0 - 0.001 '***', 0.001 - 0.01 '**', 0.01 - 0.05 '*', 0.05 - 0.1 '.', 0.1 - 1.0 (No symbol)

1w - 1 week of life, 1m - 1 month, 6m - 6 months, 1y - 1 year, 2y - 2 years.

**Supplementary Figure 7.** Country-specific significant changes in genus-level abundance from static timepoints (1 week) and overlap of genus-level signals in full cohort

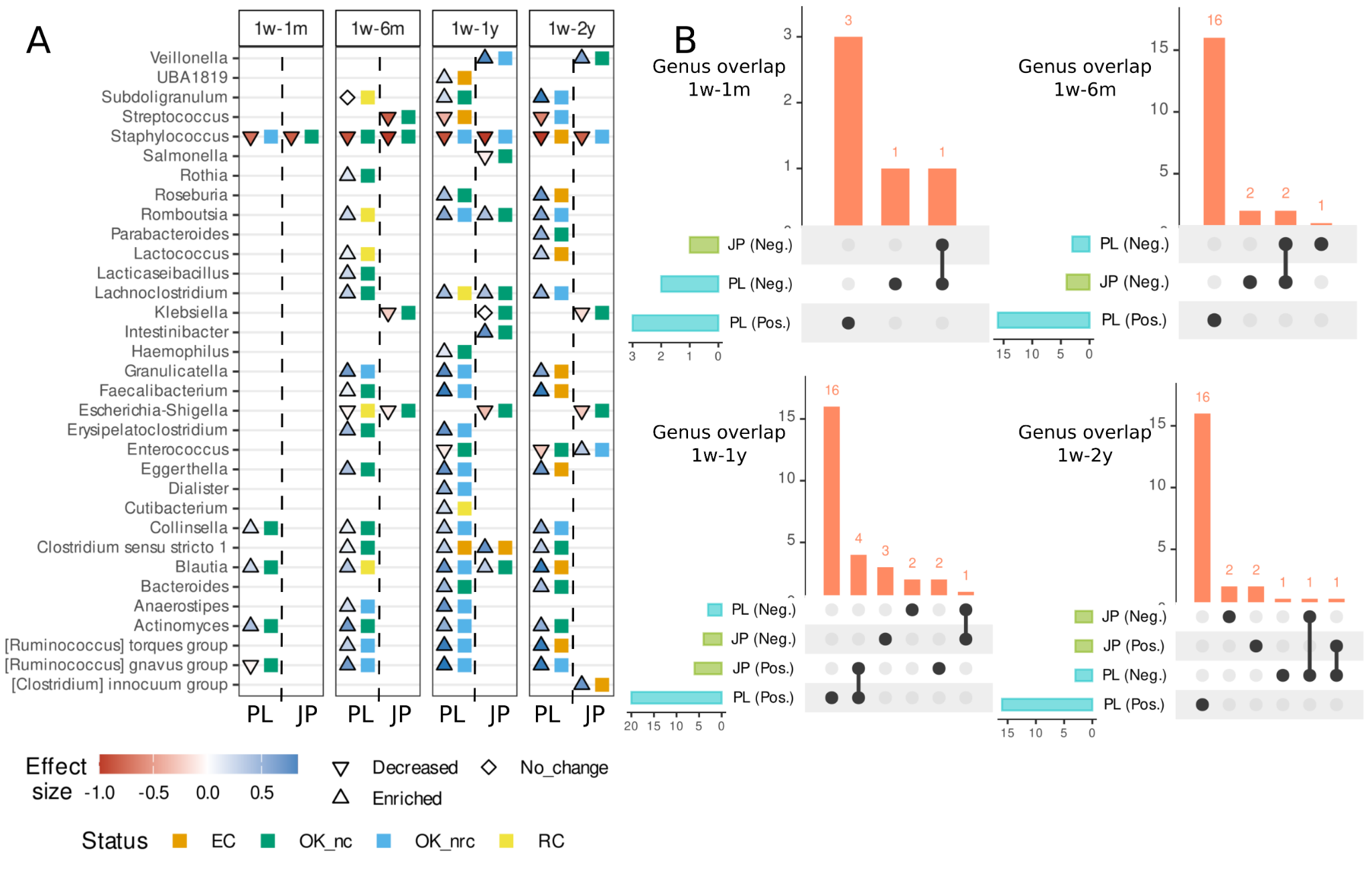

**Supplementary Figure 8**. Country-specific significant changes in genus-level abundance from static timepoints (1 month) and overlap of genus-level signals in full cohort

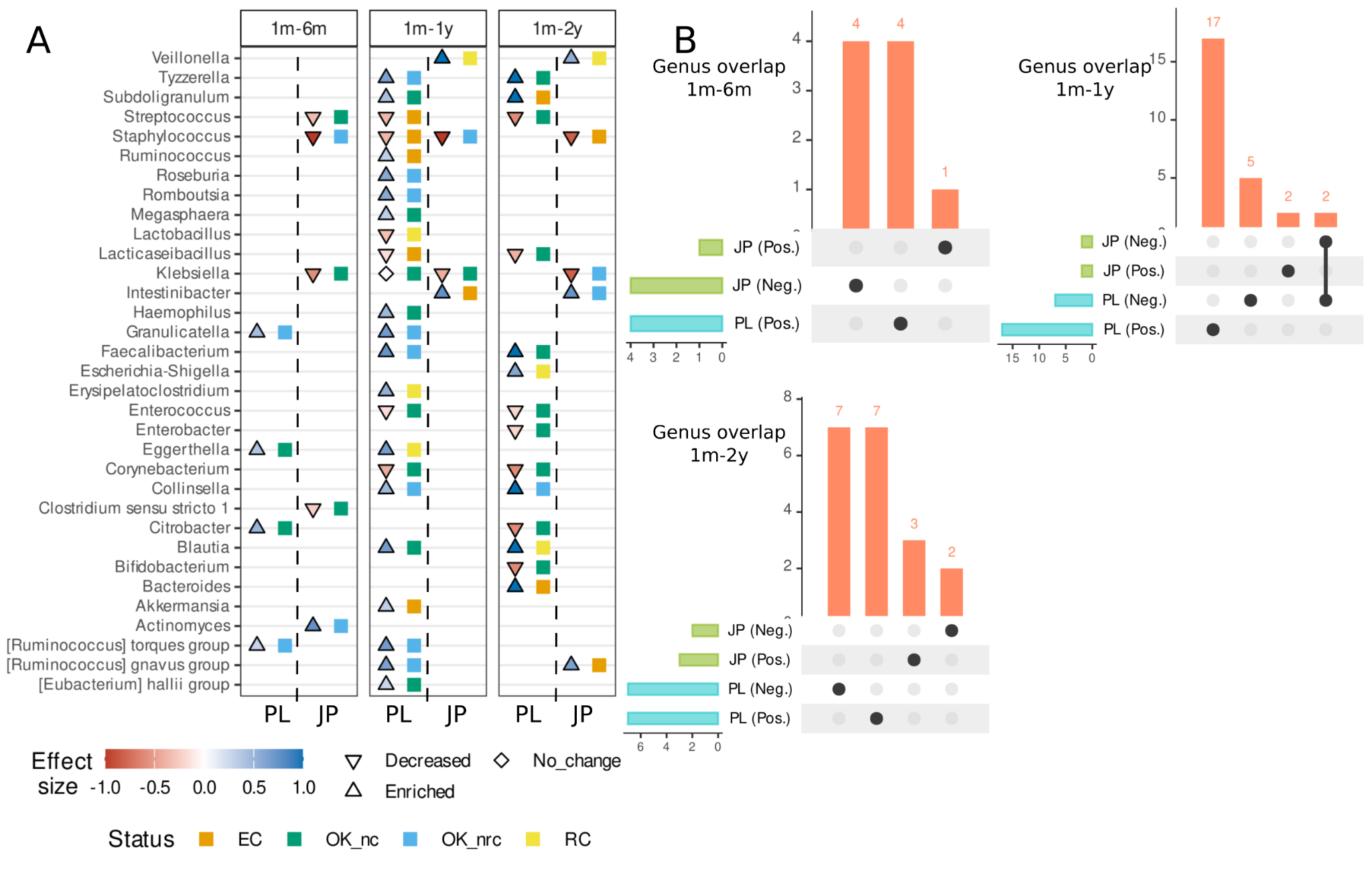

**Supplementary Figure 9**. Country-specific significant changes in genus-level abundance from static timepoints (6 months) and overlap of genus-level signals in full cohort

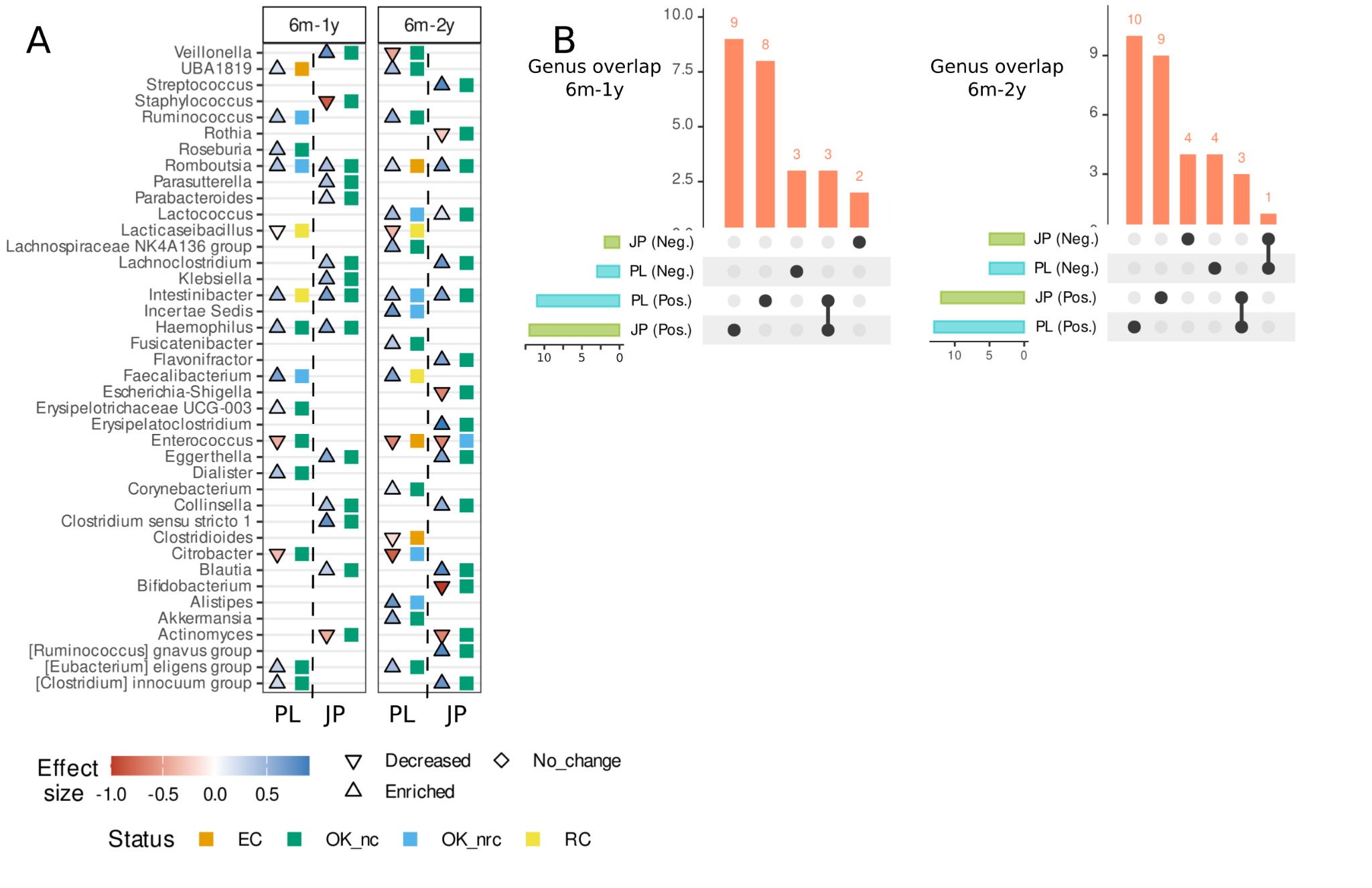

**Supplementary Figure 10**. Country-specific significant changes in genus-level abundance from static timepoints (1 week) and and overlap of genus-level signals in vaginally delivered newborns

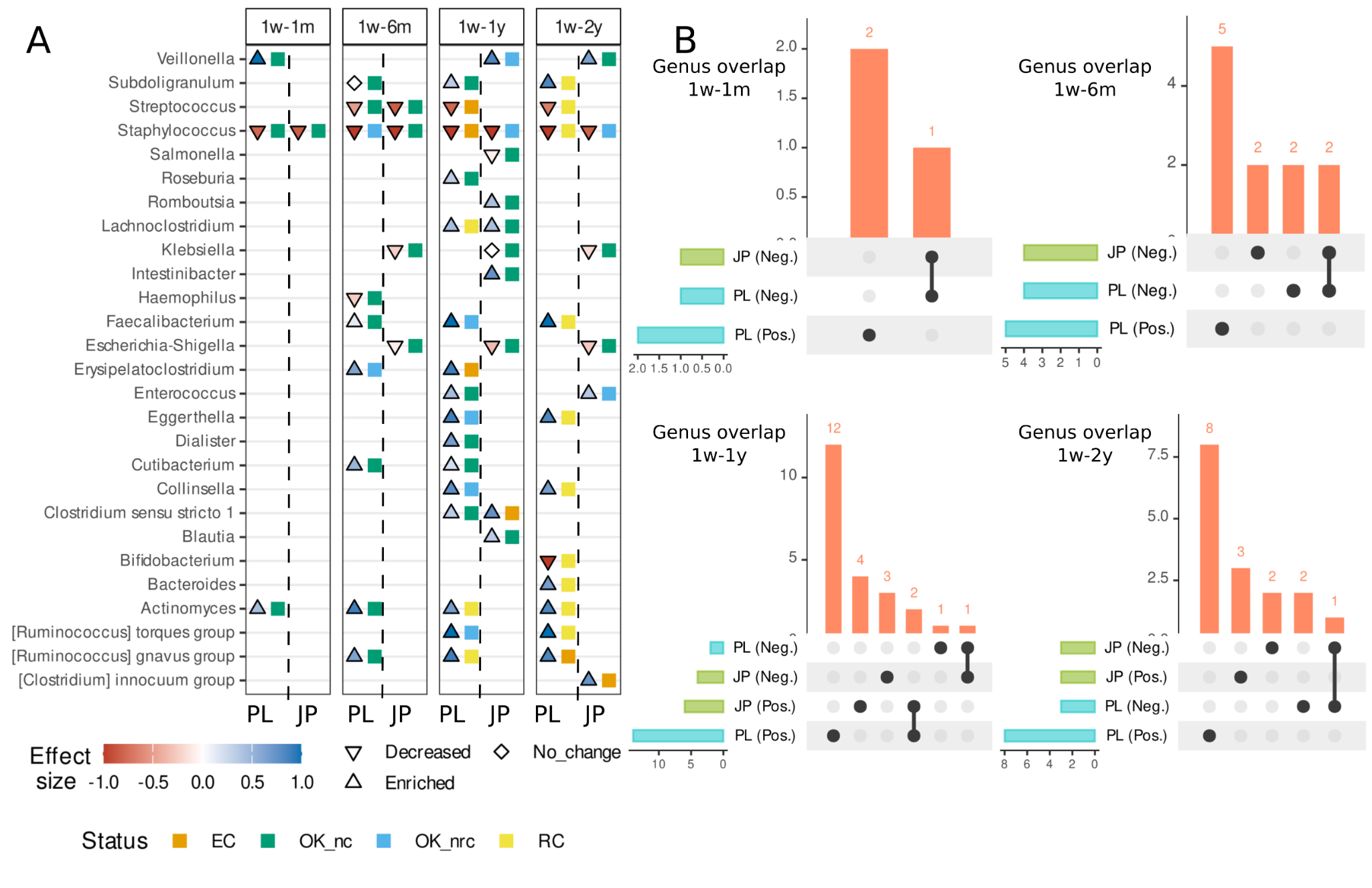

**Supplementary Figure 11**. Country-specific significant changes in genus-level abundance from static timepoints (1 month) and overlap of genus-level signals in vaginally delivered newborns

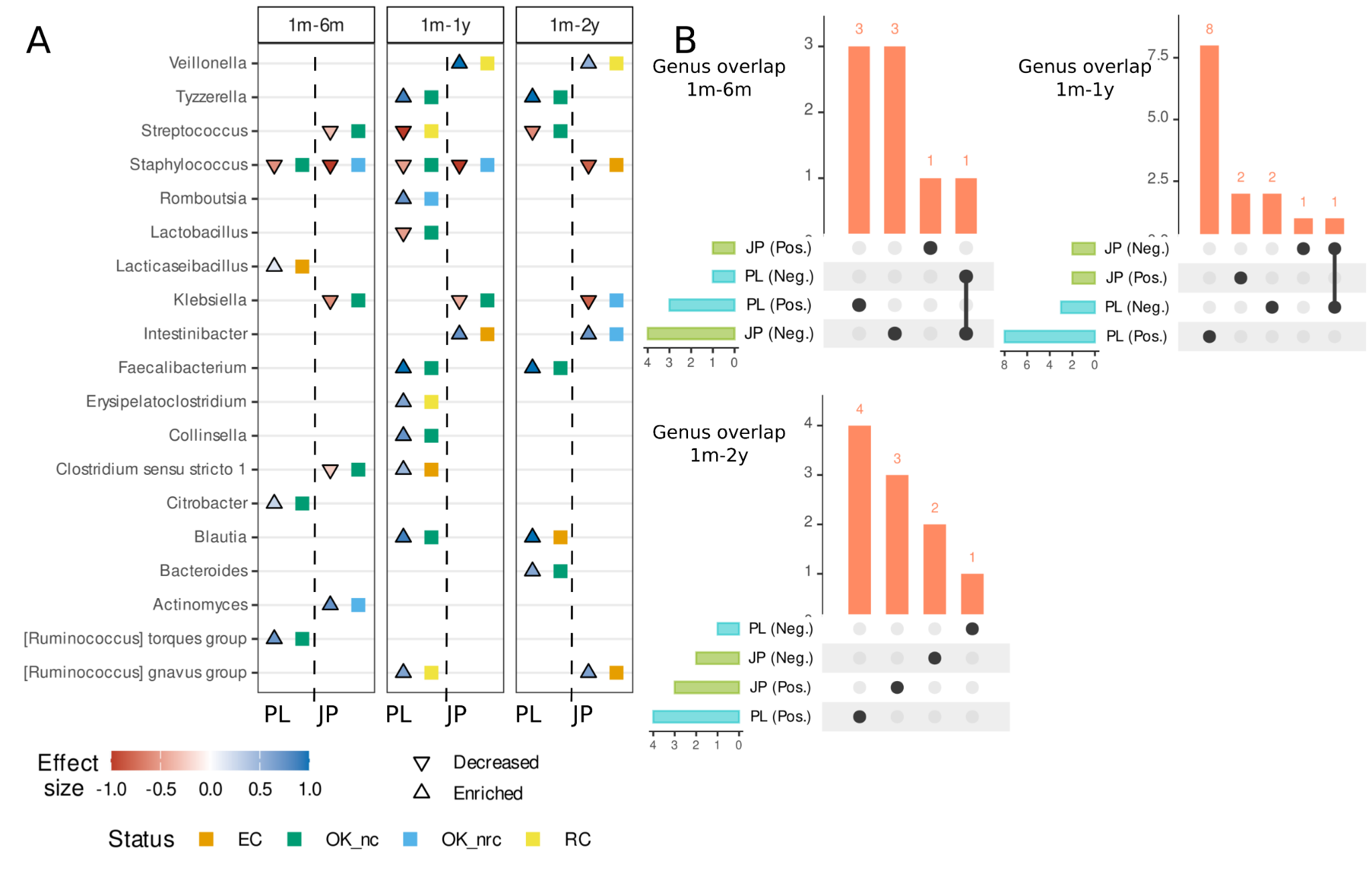

**Supplementary Figure 12**. Country-specific significant changes in genus-level abundance from static timepoints (6 months) and overlap of genus-level signals in vaginally delivered newborns

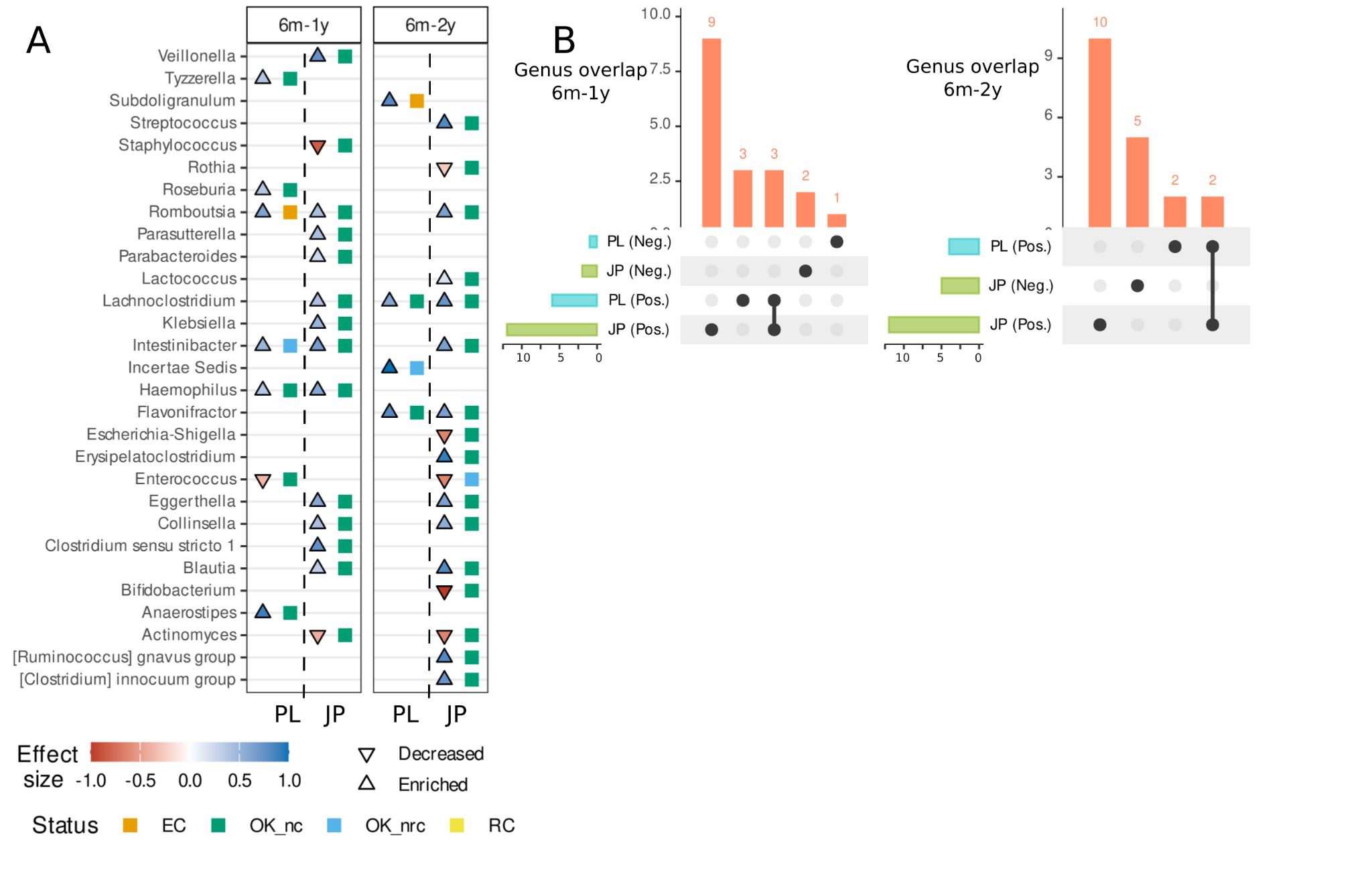

Legend for Supplementary Figures 7-12

A - cuneiform plots displaying features whose signals were not classified as “NS” (non-significant) - the direction of effects across the respective time intervals are shown. A triangle pointing downward indicates a decrease in abundance between time points, while Enriched denotes an increase. Effect sizes are represented by Cliff’s delta. Status labels definitions: EC (Entangled with covariate) - there are potential covariates, and it is not possible to conclude whether the effect is resulted from time or covariates; RC (Reducible to covariate) - there is an effect of time, but it can be reduced to the covariate effects; OK_nc (OK, no covariate) - there is an effect of time and there is no potential covariate; and OK_nrc (OK, not reducible to covariate) - there are potential covariates, but there is an effect of time and it is independent of those of covariates. B - UpSet plots illustrating the overlap of significant genera between countries. The direction of association is also indicated. The intersection size bar plot (vertical) indicates the number of genera shared across different country - effect combinations, while the set size (horizontal) bars represent the total number of genera per group: PL (Neg.) - Polish cohort, decrease in abundance between time points, PL (Pos.) - Polish cohort, increase in abundance between time points, JP (Neg.) - Japanese cohort, decrease in abundance between time points, JP (Pos.) - Japanese cohort, increase in abundance between time points; Only individuals with complete samples at the compared time points were included.

**Supplementary Figure 13**. Cohort-specific significant changes in genus-level abundance across time points (consecutive and non-consecutive) in PL, JP, FIN, and US cohorts

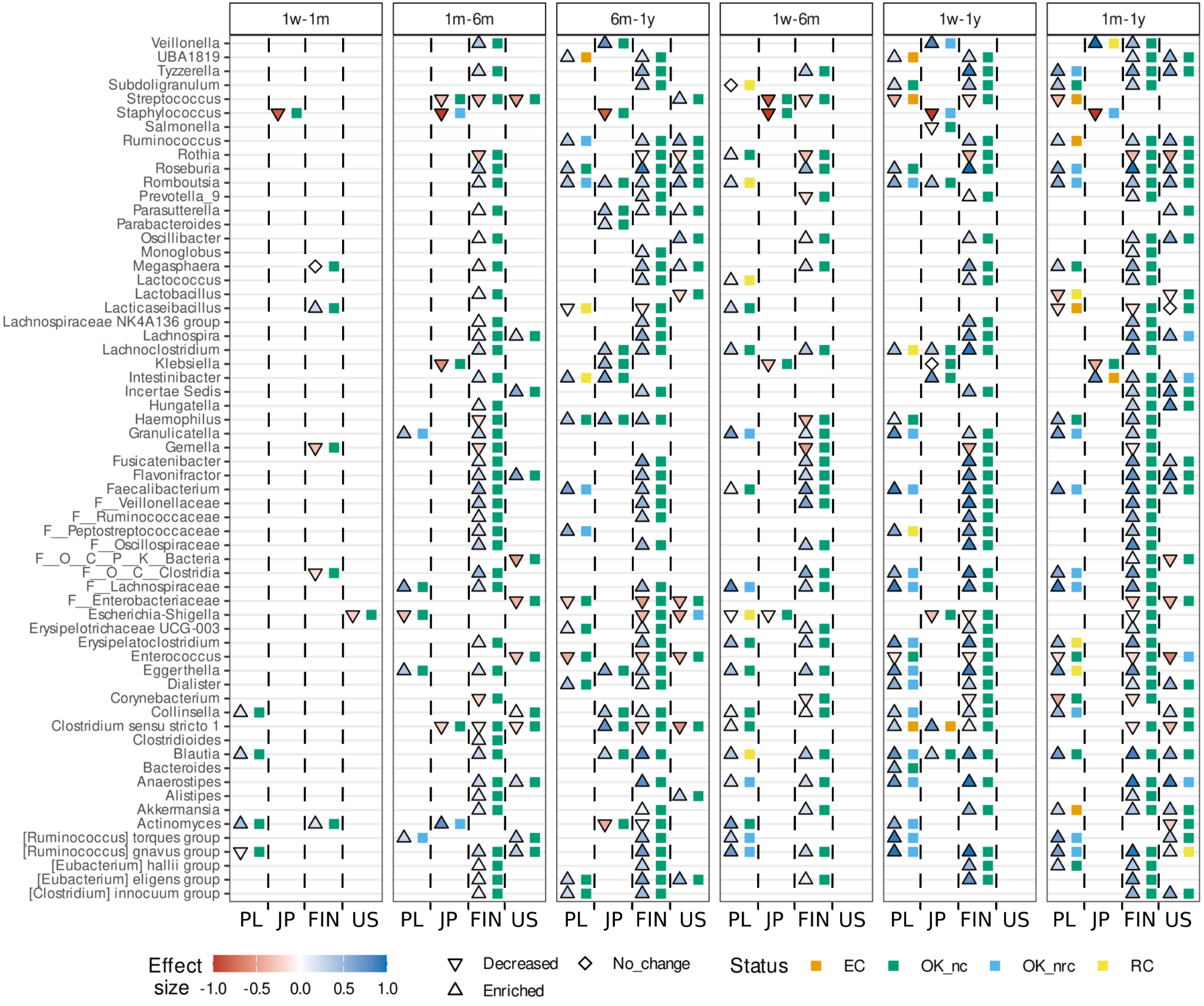

**Supplementary Figure 14**. Overlap of genus-level signals across consecutive and non-consecutive time points in PL, JP, FIN, and US full cohorts

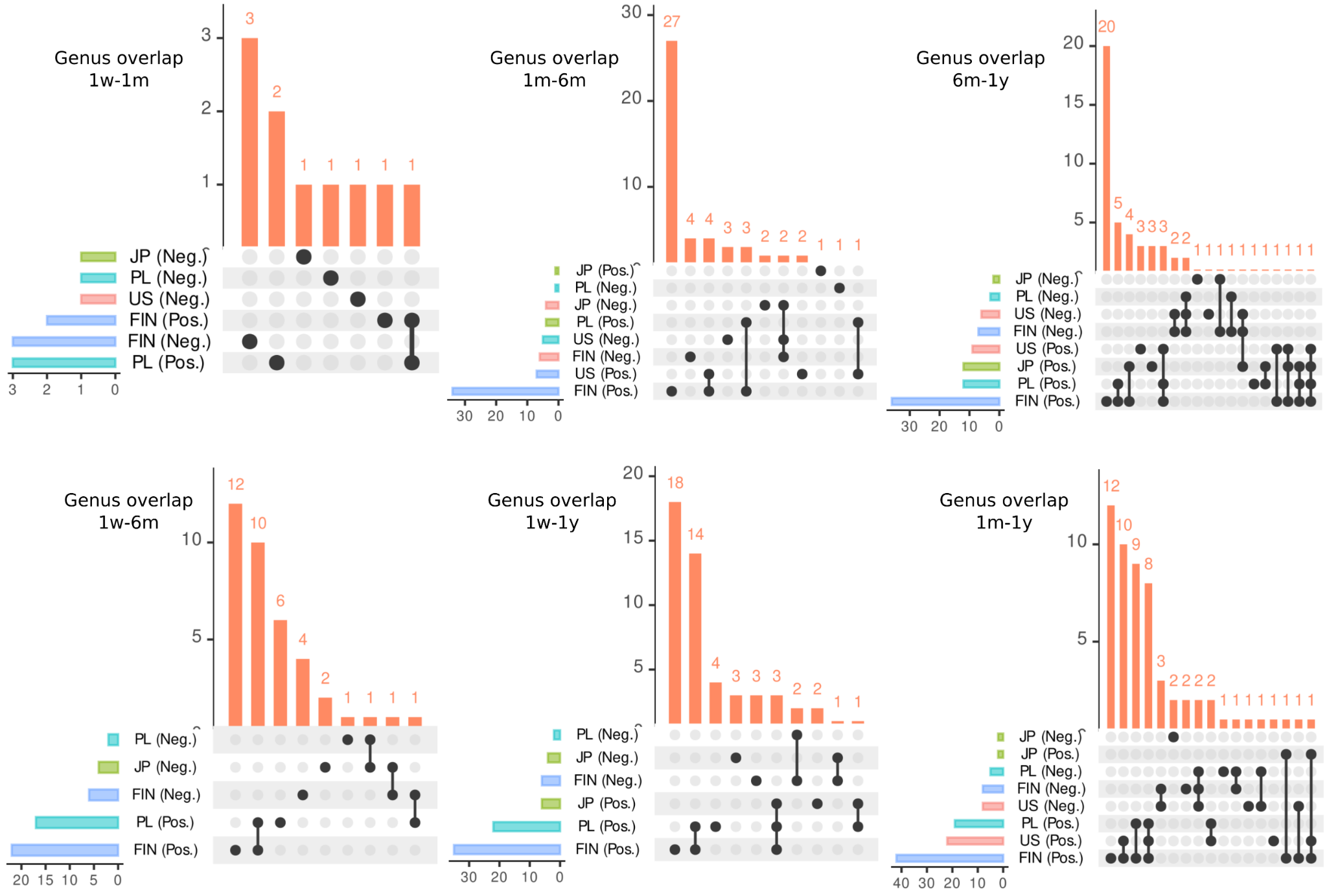

**Supplementary Figure 15**. Overlap of genus-level signals across consecutive and non-consecutive time points in PL and JP (full cohorts) and in **subsampled FIN and US cohorts**

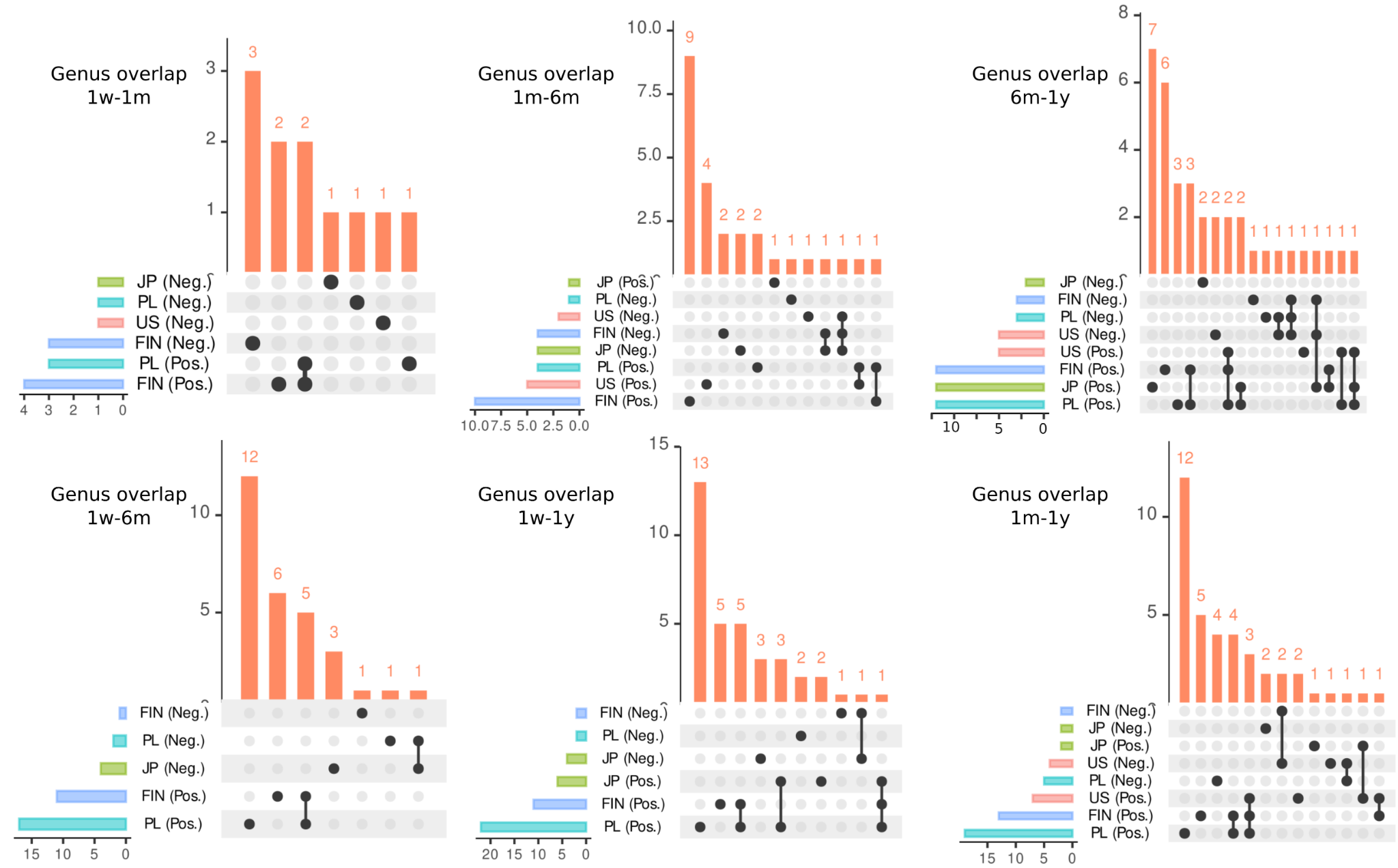

**Supplementary Figure 16.** Venn diagrams of genus-level changes across time points in subsampled FIN/US and full PL/JP cohorts

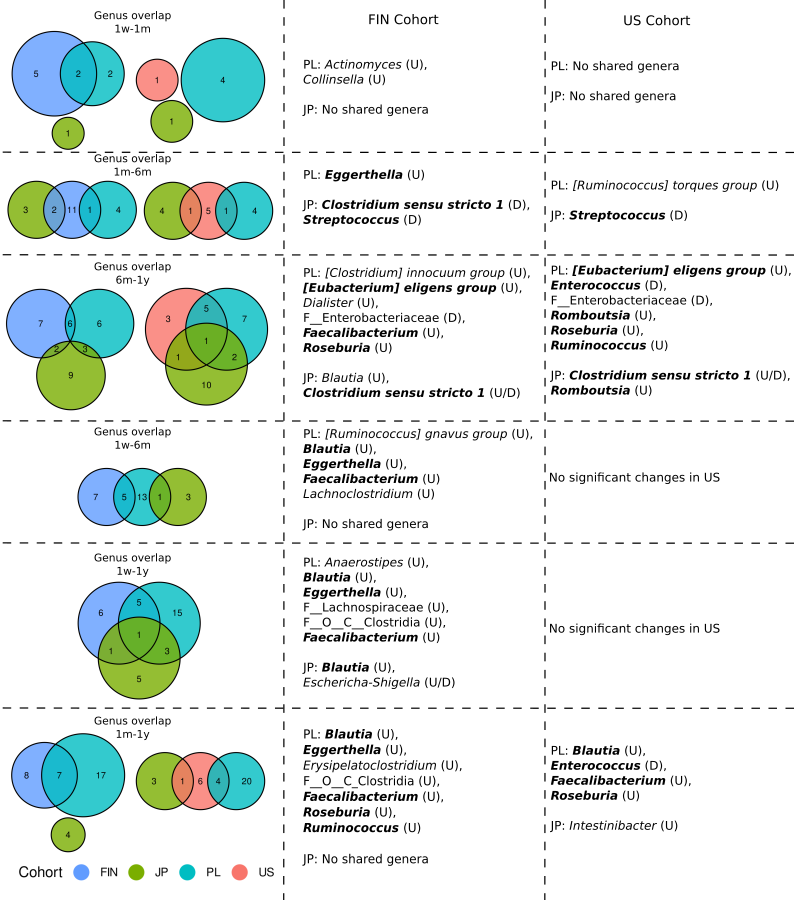

**Supplementary Tables**

**Supplementary Table 1**. Characteristics of the newborn cohorts included in the study and obtained from public resources

| Project | Study population | Study design | DNA extraction | Platform |
| --- | --- | --- | --- | --- |
| **Finnish**  **N=153 (516 samples: HELMi: 196, Jorvi: 320)** | Cohorts: Finnish Health and Early Life Microbiota (HELMi) cohort and Jorvi cohort  **Total Number of Infants: 153 (2684 stool samples)** | The primary objective of the study was to understand the impact of birth interventions (specifically, cesarean section and intrapartum antibiotics) on gut microbiota development in relation to health in early infancy. Observational longitudinal study | repeated bead-beating, DNA extraction with a KingFisher | Illumina MiSeq and HiSeq (V3V4) |
| **ECAM**  **N=38 (138 samples)** | The study population consists of 43 healthy urban infants from the United States. These infants were enrolled for follow-up for up to the age of 2 years. Additionally, stool samples were collected from these infants, and samples (stool, vaginal swabs, rectal swabs) were collected from their mothers both pre- and post-partum. | observational longitudinal cohort study focusing on the development of the intestinal microbiota in a cohort of 43 healthy urban infants over the first 2 years of life. The study aims to understand the effects of birth mode, infant nutrition, and antibiotic exposures on microbiota development in early childhood. | Bead beating;  Power-Soil-htp 96 Well Soil DNA Isolation Kit (MoBio, Carlsbad CA) | Illumina MiSeq (V4) |
| **Japanese**  **N=12 (60 samples)** | 12 healthy (1048 samples), full-term infants. All infants were Japanese, delivered vaginally, and primarily fed breast milk during the first 2 years of life. | Observational longitudinal study investigating the gut microbiome and metabolite profiles of 12 healthy full-term infants over the first 2 years of life. The design involves repeated measures and aims to explore associations among SCFA profiles and gut microbiota composition | bead-beating phenol-chloroform | Illumina MiSeq (V1V2) |

**Supplementary Table 2**. Number of newborns per time point (gut microbiota samples and metadata availability)

| Project | Birth | 1 week | 1 month | 6 months | 1 year | 2 years | Total |
| --- | --- | --- | --- | --- | --- | --- | --- |
| Before removing samples with total count <= 5,000 | | | | | | | |
| Finnish | 66 (12.8%) | 66 (12.8%) | 149 (28.9%) | 143 (27.7) | 92 (17.8%) | 0 (0%) | 516 (100%) |
| ECAM | 29 (21.0%) | 4 (2.9%) | 28 (20.3%) | 32 (23.2%) | 30 (21.7%) | 15 (10.9%) | 138 (100%) |
| Japanese | 12 (20.0%) | 12 (20.0%) | 12 (20.0%) | 12 (20.0%) | 12 (20.0%) | 12 (20.0%) | 60 (100%) |
| Polish | 10 (9.4%) | 23 (21.5%) | 20 (18.7%) | 20 (18.7%) | 21 (19.6%) | 13 (12.1%) | 107 (100%) |
| After removing samples with total count <= 5,000 | | | | | | | |
| Finnish | 9 (2.1%) | 50 (116%) | 141 (32.7%) | 141 (32.7%) | 90 (20.9%) | 0 (0%) | 431 (100%) |
| ECAM | 24 (18.2%) | 4 (3.0%) | 28 (21.2%) | 31 (23.5%) | 30 (22.7%) | 15 (11.4%) | 132 (100%) |
| Japanese | 12 (20.3%) | 12 (20.3%) | 12 (20.3%) | 12 (20.3%) | 12 (20.3%) | 11 (18.6%) | 59 (100%) |
| Polish | 9 (8.6%) | 23 (21.9%) | 20 (19.0%) | 20 (19.0%) | 21 (20.0%) | 12 (11.4%) | 105 (100%) |
|  | Sex (F/M) | Delivery (VB/CS) | ATB (Yes/No) | Diet (BM/Other) |  |  |  |
| Finnish | NA | 310/121  (71.9%/28.1%) | 248/183  (57.5%/42.5%) | NA |  |  |  |
| ECAM | 46/86  (34.8/65.2) | 77/55  (58.3%/41.7%) | 32/100  (24.2%/75.8%) | NA |  |  |  |
| Japanese | 25/34  (42.4%/57.6%) | 59  (100%) | 2/57  (3.4%/96.6%) | 19/40  (32.2%/67.8%) |  |  |  |
| Polish | 51/54  (48.6%/51.4%) | 47/58  (44.8%/55.2%) | 31/74  (29.5%/70.5%) | 59/46  (56.2%/43.8%) |  |  |  |

**Supplementary Table 3. Japanese cohort metadata for all samples (1,048) and samples included in this study**

Provided as an Excel file

**Supplementary Table 4.**  Detailed statistics* on ASVs per sample and per feature by project

| Characteristic | Finnish** | | ECAM | Japanese | Polish |
| --- | --- | --- | --- | --- | --- |
|  | MiSeq | HiSeq |  |  |  |
| Samples | 368 | 149 | 138 | 60 | 107 |
| 16S region | V3-V4 | V3-V4 | V4 | V1-V2 | V3-V4 |
| Sequence length: overall/post-filtering | 250/170 | 200/170 | 150/150 | 250/240 | 250/233 |
| Features (ASV) overall/post-filtering | 2011/1686 | 970/547 | 1649/1381 | 2166/1655 | 65676/4655 |
| Features per sample | 4.58 | 3.67 | 9.94 | 27.58 | 43.51 |
| Frequency of assigned reads per sample | | | | | |
| Minimum | 8 | 0 | 2,711 | 4,429 | 354 |
| 1st quartile | 14,624.25 | 380 | 15,052 | 10,823.5 | 82,296 |
| Median | 25,098.5 | 19,154 | 19,362.5 | 18,144 | 94,150 |
| 3rd quartile | 36,472 | 47,085 | 23,444.25 | 41,940.25 | 129,031 |
| Maximum | 110,125 | 180,773 | 66,462 | 80,206 | 635,185 |
| Mean | 26,054.88 | 31,215.87 | 19,873.8 | 25,268.05 | 111,281.5 |
| Frequency of assigned reads per feature | | | | | |
| Minimum | 4 | 3 | 3 | 11 | 5 |
| 1st quartile | 95 | 98 | 44 | 82 | 100 |
| Median | 351.5 | 413 | 80 | 139 | 281 |
| 3rd quartile | 1816.5 | 3237.5 | 201 | 557 | 1,046 |
| Maximum | 310,366 | 295,550 | 259,557 | 19,267 | 217,183 |
| Mean | 5,686.95 | 8,503.04 | 1,985.94 | 916.06 | 2,557.92 |

*before removing samples with total count <= **5,000** (See Supplementary methods), ** 517 microbiota samples, 516 metadata available

**Supplementary Table 5. Cross-sectional comparisons between the PL and JP cohorts, covariate-adjusted, each time point - non-restricted**

Provided as an Excel file

**Supplementary Table 6. Cross-sectional comparisons between the PL and JP cohorts, covariate-adjusted, each time point - restricted**

Provided as an Excel file

**Supplementary Table 7**. **Paired non-resticted vs restricted (ASVs mapped to shared genera) comparison for each cohort and time point**

Provided as an Excel file

**Supplementary Table 8**. PERMANOVA results across all scenarios (full set, shared genera, unadjusted, and covariate-adjusted) - Polish vs Japanese cohorts

Provided as an Excel file

**Supplementary Table 9**. Differential abundance analysis - confounder-aware approach (using the metaDeconfoundR R package)

Provided as an Excel file

**Supplementary Table 10**. **Cross-sectional comparisons between the PL and JP cohorts, covariate-adjusted, each time point - non-restricted, vaginally born only**

Provided as an Excel file

**Supplementary Table 11. Cross-sectional comparisons between the PL and JP cohorts, covariate-adjusted, each time point - restricted, vaginally born only**

Provided as an Excel file

**Supplementary Table 12**. **Paired non-restricted vs restricted (ASVs mapped to shared genera) comparison for each cohort and time point, vaginally born only**

Provided as an Excel file

**Supplementary Table 13**. PERMANOVA results across all scenarios (full set, shared genera, unadjusted, and covariate-adjusted) - Polish vs Japanese cohorts, vaginally-born

Provided as an Excel file

**Supplementary Table 14**. Differential abundance analysis in vaginally-born newborns - confounder-aware approach (using the metaDeconfoundR R package)

Provided as an Excel file

**Supplementary Table 15**. Overlap of genus-level signals in full and vaginally-born cohorts

Provided as an Excel file

**Supplementary Table 16**. Metadata of all four cohorts included in the study

Provided as an Excel file
